# The FDA-approved drug cobicistat synergizes with remdesivir to inhibit SARS-CoV-2 replication

**DOI:** 10.1101/2021.03.09.434219

**Authors:** Iart Luca Shytaj, Mohamed Fares, Bojana Lucic, Lara Gallucci, Mahmoud M. Tolba, Liv Zimmermann, Ahmed Taha Ayoub, Mirko Cortese, Christopher J. Neufeldt, Vibor Laketa, Petr Chlanda, Oliver T. Fackler, Steeve Boulant, Ralf Bartenschlager, Megan Stanifer, Andrea Savarino, Marina Lusic

## Abstract

Combinations of direct-acting antivirals are needed to minimize drug-resistance mutations and stably suppress replication of RNA viruses. Currently, there are limited therapeutic options against the Severe Acute Respiratory Syndrome Corona Virus 2 (SARS-CoV-2) and testing of a number of drug regimens has led to conflicting results. Here we show that cobicistat, which is an-FDA approved drug-booster that blocks the activity of the drug metabolizing proteins Cytochrome P450-3As (CYP3As) and P-glycoprotein (P-gp), inhibits SARS-CoV-2 replication. Cell-to-cell membrane fusion assays indicated that the antiviral effect of cobicistat is exerted through inhibition of spike protein-mediated membrane fusion. In line with this, incubation with low micromolar concentrations of cobicistat decreased viral replication in three different cell lines including cells of lung and gut origin. When cobicistat was used in combination with the putative CYP3A target and nucleoside analog remdesivir, a synergistic effect on the inhibition of viral replication was observed in cell lines and in a primary human colon organoid. The cobicistat/remdesivir combination was able to potently abate viral replication to levels comparable to mock-infected cells leading to an almost complete rescue of infected cell viability. These data highlight cobicistat as a therapeutic candidate for treating SARS-CoV-2 infection and as a potential building block of combination therapies for COVID-19.

## Introduction

The ongoing pandemic of Severe Acute Respiratory Syndrome Corona Virus type 2 (SARS-CoV-2) poses the challenge of quick development of antiviral therapies. SARS-CoV-2 is an enveloped, positive sense, RNA virus of the *Coronaviridae* family, which includes other human-infecting pathogens such as SARS-CoV and MERS-CoV (1). Currently, there are no widely approved antivirals to treat infection with Coronaviruses. Substantial effort has been devoted to identifying inhibitors of SARS-CoV-2 replication through repurposing of compounds approved for treating other clinical indications. Repositioned drugs offer the advantage of a well-known safety profile and the possibility of faster clinical testing, which is essential during a sudden epidemic outbreak (2). Large scale clinical trials have identified immune modulating agents (*e*.*g*. dexamethasone (3,4) and baricitinib (5)) as potential treatments for Coronavirus disease 2019 (COVID-19). However, direct acting antiviral agents have shown limited clinical benefits so far. In particular, a set of antiviral drugs initially identified as effective *in vitro* (remdesivir, chloroquine/hydroxychloroquine) has been unable to reproducibly decrease mortality in placebo controlled trials (6–9).

Complete inhibition of SARS-CoV-2 replication will likely require combinations of antivirals, in line with previous evidence on other RNA viruses (10,11). Candidate inhibitors have been proposed to target several critical steps of SARS-CoV-2 replication, including viral entry, polyprotein cleavage by viral proteases, transcription and viral RNA replication (12). SARS-CoV-2 entry is mediated by the spike glycoprotein (S-glycoprotein), which binds through its S1 subunit to the cellular receptor Angiotensin-converting enzyme 2 (ACE2). Upon binding, the viral entry requires a proteolytic activation of the S2 subunit leading to the fusion of the viral envelope with the host cell membrane (13). The study of candidate inhibitors of SARS-CoV-2 entry has mainly focused on monoclonal antibodies and small molecules to target the association of the receptor binding domain (RBD) of the S-glycoprotein to ACE-2 (14). Interestingly, the intensively studied antimalarials chloroquine and hydroxychloroquine have been suggested to impair SARS-CoV-2 entry *in vitro* by both decreasing the binding of the RBD to ACE2 and by decreasing endosomal acidification (15,16).

Upon viral membrane fusion, the viral RNA is released to the cytosol and translated into two large polyproteins that are cleaved into non-structural proteins (nsp′s) by two viral proteases, the main protease (3CL_pro_) and the papain-like protease (PL_pro_). A large body of work to identify antivirals against SARS-CoV-2 has focused on research on these viral proteases (17). Initial drug repurposing efforts focused on inhibitors of the HIV-1 protease, such as lopinavir and darunavir, alone or in combination with pharmacological boosters (*i*.*e*. ritonavir and cobicistat). These inhibitors, however, proved poorly effective in inhibiting 3CL_pro_ activity *in vitro (18)* and did not offer reproducible clinical benefit (19–21). Larger drug screenings have so far relied on a combination of *in-silico* and *in-vitro* tools (22). In particular, libraries of compounds have been screened through molecular docking and many candidate drugs have shown favorable binding properties to the SARS-CoV-2 proteases when analyzed by molecular dynamics (23). In the laboratory, however, repurposed inhibitors of SARS-CoV-2 proteases have generally shown half-maximal inhibitory concentration (IC50) values that were incompatible with dosages achievable *in vivo*.

The nsps generated by polyprotein cleavage by the viral proteases support the transcription and replication of the viral genome, which is catalyzed by the activity of the RNA-dependent RNA polymerase (RdRP). Owing to its crucial role and high evolutionary conservation, this viral enzyme represents a very attractive therapeutic target, which has so far been exploited by repurposing the anti-Ebola virus drug remdesivir (7,8,24). Other potential RdRP inhibitors, repurposed from treatment of HCV, HIV-1 and influenza virus have been proposed as well (2 5– 27). Among them, Favipravir and Molnupiravir (MK-4482) have shown *in-vivo* therapeutic potential by decreasing viral burden and transmission in hamster and ferret models of the infection, respectively (28,29).

Viral transcripts generated by the RdRP are used for assembly of new virions by budding into the lumen of the ER-Golgi intermediate compartment (ERGIC) (30). The assembly is driven by the structural proteins M and E which are responsible for the incorporation of the N protein forming ribonucleoprotein complexes containing the viral genome. After the budding is completed, the virions are released from the cell either by exocytosis or through lysosomal organelle trafficking (1). So far, drug candidates proposed to target viral assembly/budding have not advanced beyond *in-silico* predictions (31).

A major limitation hampering the development of combined antiviral strategies against SARS-CoV-2 is the paucity of data available on drug interactions. Initial guidelines and *in-vitro* results have discouraged the combined use of potentially effective compounds, such as remdesivir and chloroquine/hydroxychloroquine, on the basis of possible antagonism (32) or interference of chloroquine/hydroxychloroquine with remdesivir metabolism through the efflux pump P-glycoprotein (P-gp) [(Gilead. Summary on compassionate use) and (33,34)]. On the other hand, extensive first pass metabolism by the liver hampers the oral bioavailability of remdesivir and circumscribes its use to intravenous administration, thus limiting both its scalability and, likely, antiviral efficacy (35). More broadly, the family of cytochrome P450 proteins (CYP) or P-gp are responsible for the breakdown and clearance of a large majority of drugs. For this reason, compounds inhibiting CYP function find extensive use in combination therapies (*e*.*g*. against HIV-1) (36–39).

Here, we demonstrate that the FDA-approved CYP3A inhibitor cobicistat, typically used as a booster of HIV-1 protease inhibitors (40), can block SARS-CoV-2 replication *in vitro* in cell lines of lung and gut origin. While cobicistat was identified through *in-silico* screening by several groups as a potential inhibitor of 3CL_pro_, our data point towards an effect on the S-protein, which, in the presence of cobicistat, showed decreased ability to form syncytia in cells overexpressing the S-protein. The antiviral concentrations of cobicistat, while well tolerated *in vitro*, are above those typically used for HIV-1 treatment, but compatible with plasma levels previously reached at higher doses in mice as well as in humans. In combination with remdesivir, cobicistat exhibits a synergistic effect in rescuing cell viability and abrogating viral replication in both cell lines and in a primary colon organoid. Overall, our data show that cobicistat has a dual activity both as an antiviral drug and as a pharmacoenhancer, thus potentially constituting a basis for combined therapies aimed at complete suppression SARS-CoV-2 replication.

## Results

### *In-silico* and *in-vitro* analyses identify cobicistat as a candidate inhibitor of SARS-CoV-2 replication

To identify potential inhibitors of SARS-CoV-2 replication we performed a structure-based virtual screening of the Drugbank library of compounds approved for clinical use. Candidate drugs were ranked based on their docking score to the substrate-binding site of 3CL_pro_, *i*.*e*. the site essential for the proteolytic function. Our results highlighted seventeen top candidate inhibitors, including compounds used to treat parasitic as well as viral infections (Supplementary Table 1). Among the latter, the HIV-1 protease inhibitor nelfinavir, which was one of the top scoring compounds in our analysis, was previously shown to decrease SARS-CoV and SARS-CoV-2 replication *in vitro (41,42)* (Supplementary Table 1). Two additional drugs used for treatment of HIV-1 displayed top docking scores, *i*.*e*. the protease inhibitor tipranavir and the CYP3A inhibitor cobicistat. The latter was a particularly interesting candidate, given its activity as a booster for HIV-1 protease inhibitors (40), which renders it a promising candidate for combination therapies. Additional *in-silico* investigation of the binding poses of cobicistat to the 3CL_pro_ of SARS-CoV-2 corroborated the potential affinity of this drug 3CL_pro_ (Figure 1A,B). Moreover, our results were in line with similar independent analyses of other groups (43–45)

**Figure 1.**
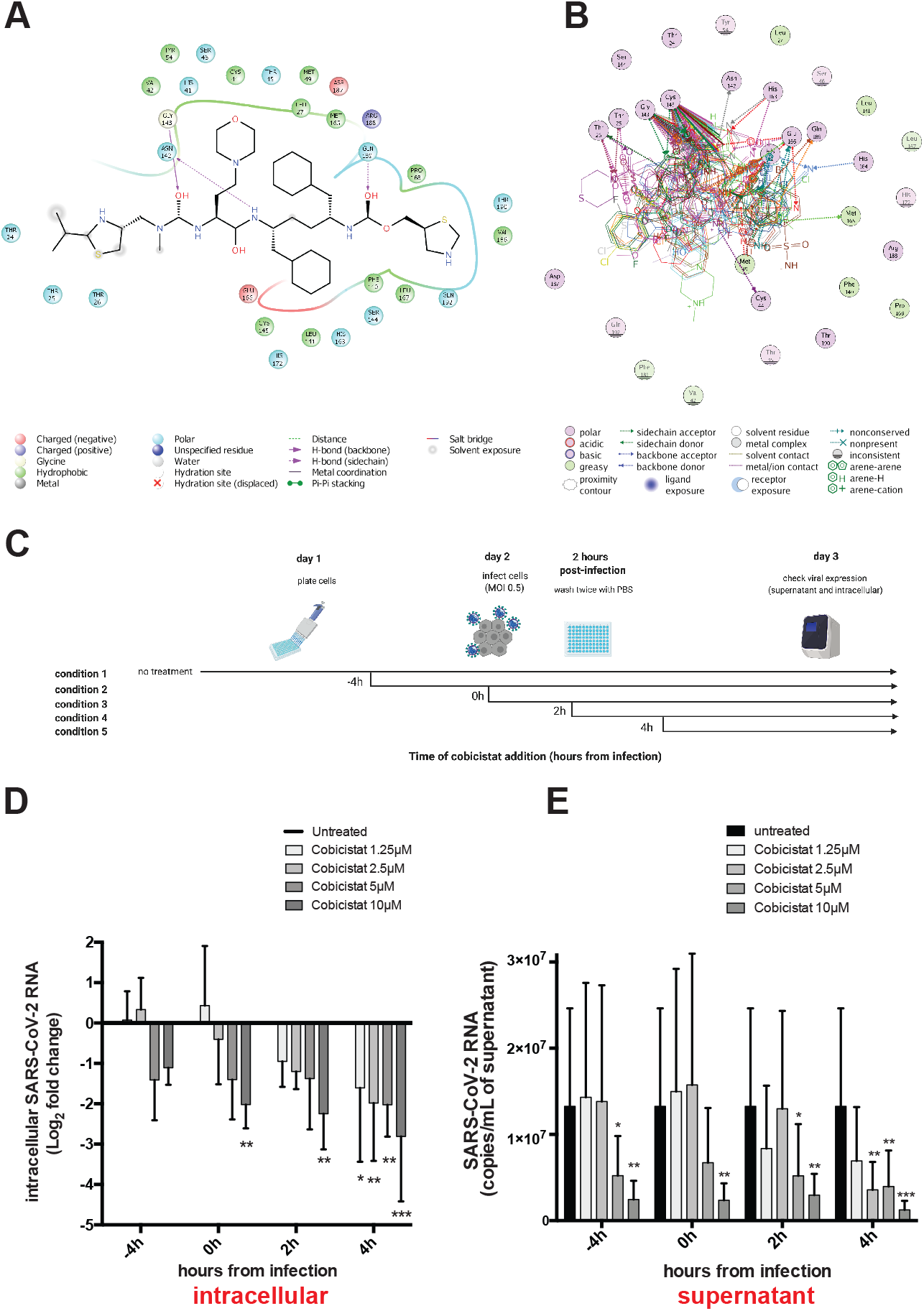
Cobicistat is a candidate inhibitor of SARS-CoV-2 replication. Panels A,B) *In-silico* docking analysis of the putative mode and energy of binding of cobicistat to SARS-CoV-2 3CL_pro_. A) Docking pose showing the ligand interaction of cobicistat to the active site of 3CL_pro_ and the formation of hydrogen bonds to ASN142, GLY143 and GLN189 of 3CL_pro_. B) Overlay of crystal structures of SARS-Cov-2 3CL_pro_ showing the amino acids important for the binding of cobicistat to the active site of the enzyme. Residues of the catalytic dyad (Cys145 and His41) of 3CL_pro_ were among the highest contributors to non covalent binding to cobicistat. The source and list of structures used are detailed in the Methods section. Panel C) Schematic representation of time course experiments evaluating *in-vitro* inhibition of SARS-CoV-2 replication by cobicistat (created with BioRender). Panels D,E) Effect of various concentrations of cobicistat, added according to the scheme of panel D, on intracellular and supernatant SARS-CoV-2 RNA content in Calu-3 cells. Viral RNA content was measured by qPCR using the 2019-nCoV_N1 primer set (Center of Disease Control). Fold change values in intracellular RNA (D) were calculated by the delta-delta CT method(82), using the Tata-binding protein (*TBP*) gene as housekeeper control. Expression levels in supernatant (E) were quantified using an *in-vitro* transcribed standard curve generated as described in the Methods section. Data are expressed as mean with SD and were analyzed by two-way ANOVA followed by Dunnet′s post-test (N = 3 independent experiments). * P < 0.05; ** P < 0.01; *** P < 0.001.

We then tested the effect of cobicistat on SARS-CoV-2 replication *in vitro*. For this purpose, we conducted a time course analysis of the effect of different concentrations of cobicistat on intracellular viral RNA replication and release of virus into the culture supernatant of Calu-3 cells (Figure 1C-E; Supplementary Figure 1A,B). Analysis of virus RNA amounts by qPCR showed a dose dependent inhibitory effect of low micromolar concentrations of cobicistat (Figure 1D,E; Supplementary Figure 1A,B). This effect was visible in both supernatants and cellular extracts, and was reproducible when samples were assayed with two different sets of primers [*i*.*e*. N1 and N2 primer sets (Supplementary Table 2) recommended by the Center of Disease Control (Figure 1D,E; Supplementary Figure 1A,B)]. Of note, pre-incubation or treatment upon infection with cobicistat did not increase the antiviral effects as compared to adding the drug two or four hours post-infection, potentially suggesting an effect on late stages of the viral life cycle.

Taken together, these data show that cobicistat has a direct antiviral effect on SARS-CoV-2 replication *in vitro*.

### The antiviral concentration range of cobicistat is well tolerated *in vitro* and compatible with plasma levels achievable in humans and mice

We next analyzed more thoroughly the antiviral effects of cobicistat using three cell lines of different origin, *i*.*e*. Calu-3 cells (human lung), Vero E6 cells (african green monkey kidney) and T84 cells (human gut), to reflect various known or putative tissue compartments of SARS-CoV-2 replication. Each cell line was infected using two different multiplicities of infection [(MOI) 0.05 and 0.5] and left untreated or treated with various concentrations of cobicistat 2h post-infection. In all cell lines, cobicistat showed a dose dependent effect in decreasing viral RNA release in supernatant (Figure 2A). In line with this, the higher concentrations of cobicistat tested (5-10μM) were able to partially rescue viability of infected cells, as shown by both MTT and crystal violet assay (Figure 2A,B), while being well tolerated by uninfected cells (Figure 2A, Supplementary Figure 2). Overall, the range of IC50 concentrations of cobicistat (0.58-8.76μM) was dependent on the MOI of the infection and on the cell type, but always far below the half cytotoxic concentration (CC50) range of the drug on the same cell types (38.66-53μM). We then compared our *in-vitro* results with previously known pharmacokinetic properties of cobicistat in humans and mice. Interestingly, maximum plasma concentrations achievable through standard dosing of cobicistat (150mg/day as a booster for HIV-1 protease inhibitors) (37) were well below (≈1μM) most IC50 values obtained in our experiments (Figure 2C). This result is in line with the lack of effect, or only partial benefit, reported when cobicistat-boosted darunavir was tested as treatment of SARS-CoV-2 patients (20,21). On the other hand, plasma levels achievable through a higher dosage of cobicistat, which was well tolerated in clinical trials (400mg/day) (46), were above IC50 values calculated when cells were infected using a 0.05 MOI (Figure 2C). Moreover, plasma levels achievable in mice through a higher cobicistat dosage shown to be safe in this animal model (50mg/Kg) were clearly above all IC50 values calculated in our experiments, while remaining below the CC50 concentrations (Figure 2C).

**Figure 2.**
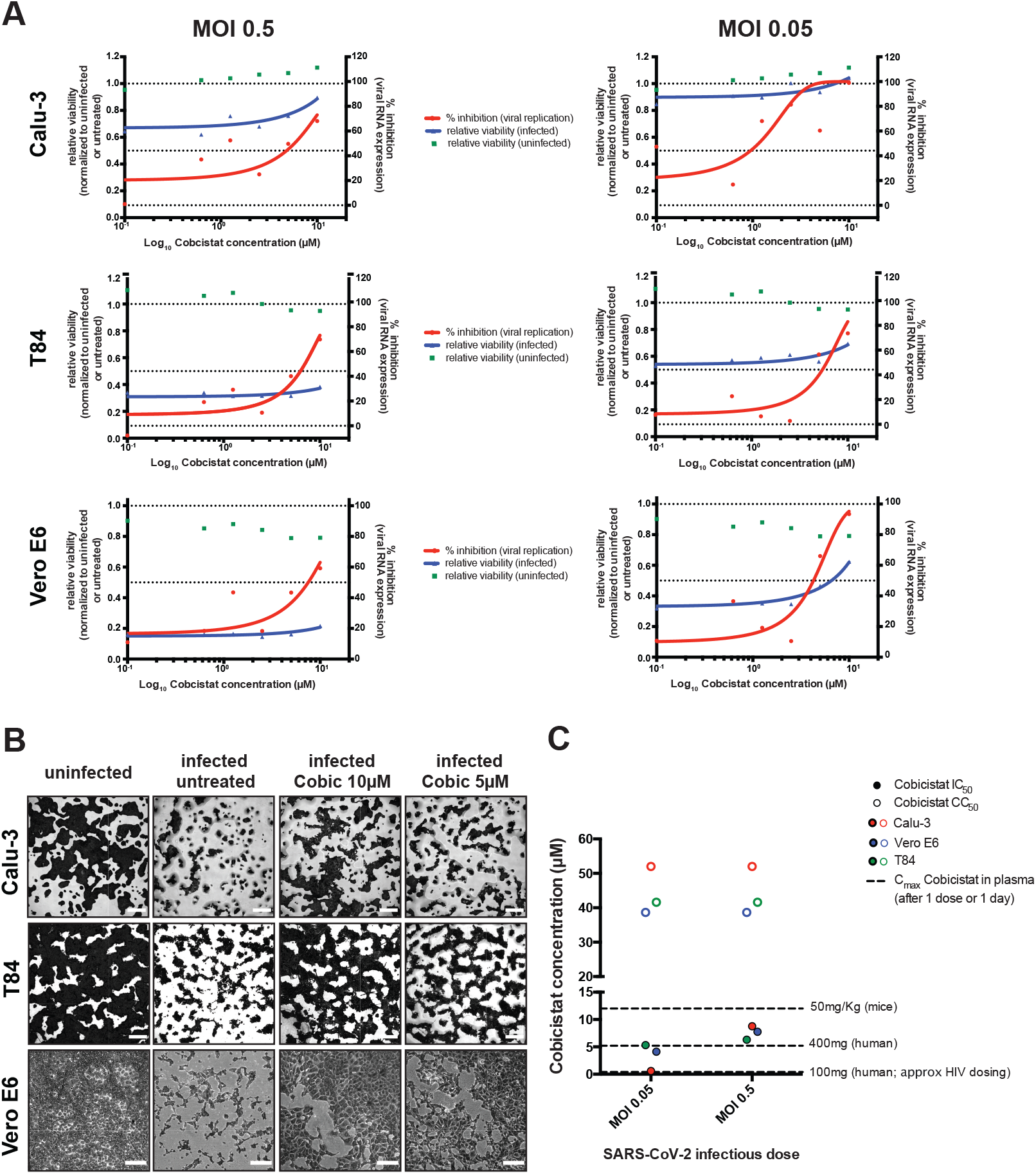
Cobicistat decreases replication of SARS-CoV-2 and rescues viability of infected cells in multiple *in-vitro* models. Panels A,B) Effect of serial dilutions of cobicistat on SARS-CoV-2 RNA concentration in supernatants (A) and on the viability of infected and uninfected cell lines of lung (Calu-3), gut (T84) and kidney (Vero E6) origin (A,B). Cells were infected with SARS-CoV-2 at two different MOIs (0.05 and 0.5) and left untreated or treated with cobicistat two hours post-infection. Forty-eight hours post-infection supernatants were collected and viral RNA was assayed by qPCR while cellular viability was measured by MTT assay (A) or by crystal violet staining (B). Inhibition of viral replication was calculated as described in the Methods section while viability data were normalized to the uninfected or to the untreated control. Half maximal inhibitory (IC50) concentration values were calculated by nonlinear regression. Each point in panel A represents a mean of 3 independent experiments. Pictures in panel B are derived from infections at MOI 0.5 (Calu-3 and T84 cells) or MOI 0.05 (Vero E6 cells). Panel C) Comparison between the IC50 and CC50 values of cobicistat determined *in vitro* and the peak plasma levels detectable in mice (Pharmacology Review of Cobicistat - Application number: 203-094) and in humans (46,88) after administration of a single dose of the drug. Determination of *in-vitro* CC50 values is based on the data shown in Supplementary Figure 2.

Overall, our data show that non-toxic concentrations of cobicistat can consistently decrease SARS-CoV-2 replication in various cellular infection models. Moreover, these data suggest that higher doses of cobicistat, as compared to the standard of care for HIV-1/AIDS, appear to be required to achieve plasma levels within the concentration range predicted to display antiviral activity.

### Cobicistat decreases S-glycoprotein content and syncytia formation *in vitro*

To characterize the mechanism of the antiviral effects of cobicistat, we analyzed the catalytic activity of 3CL_pro_ using a previously described Fluorescence Resonance Energy Transfer (FRET) assay (47). While treatment with known inhibitors of 3CL_pro_, such as GC376 and MG-132, potently reduced the catalytic activity of the enzyme, cobicistat was surprisingly inactive (Figure 3A,B). Among the top scoring compounds in our docking analysis (Supplementary Table 1), only tipranavir proved able to partially inhibit 3CL_pro_ activity, although at relatively high concentrations [half maximal effective concentration (EC50) 47μM; Figure 3A, Supplementary Table 3]. A lack of binding stability of the ligands within the active site of 3CL_pro_ might explain the discrepancy between the FRET experimental results and the previous *in-silico* predictions indicating cobicistat as a 3CL_pro_ inhibitor. In particular, when assessment of conformational entropy was included in the molecular dynamics analysis(48), the binding of cobicistat tended to be less stable than that of other ligands (Supplementary Table 3; Supplementary Movie 1) and the predicted binding energies to 3CL_pro_ of the examined ligands reflected more closely the EC50 values calculated by FRET (Supplementary Figure 3; Supplementary Table 3).

**Figure 3.**
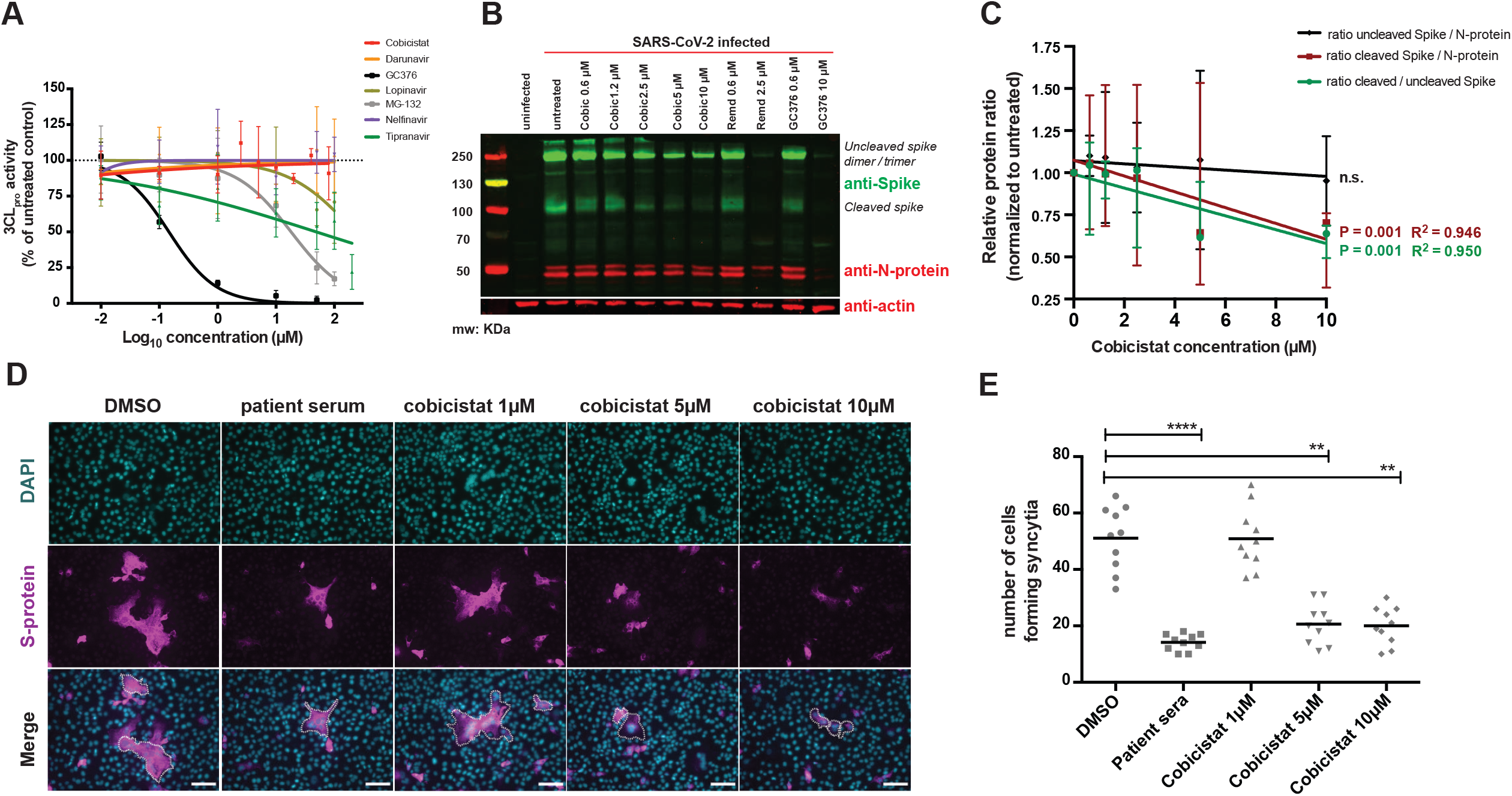
Cobicistat decreases SARS-CoV-2 S-protein content and syncytia formation. Panel A) Screening of putative inhibitors of the enzymatic activity of 3CL_pro_. The activity of 3CL_pro_ was measured by FRET assay (47) and normalized over the untreated condition. Apart from cobicistat, compounds tested included HIV-1 protease inhibitors highlighted by our molecular docking [Nelfinavir, Tipranavir (Supplementary Table 1)] or previously administered in clinical trials as SARS-CoV-2 therapeutics [darunavir (20), lopinavir (19)], as well as two positive controls known to inhibit 3CL_pro_ activity [MG-132 and GC376 (89)]. Panels B,C) Effect of cobicistat on the expression of S- and N-proteins in SARS-CoV-2 infected Vero E6 cells. Cells were infected at 0.5 MOI and left untreated or treated, two hours post-infection, with various concentrations of cobicistat, of the RdRP inhibitor remdesivir, or the 3CL_pro_ inhibitor GC376. Cells were harvested 24 hours post-treatment and subjected to protein extraction and subsequent analysis by Western Blot. Expression of S- and N-proteins, and expression of the housekeeping protein actin-β, were detected using primary monoclonal antibodies followed by incubation with fluorescent-conjugated secondary antibodies and detection on a LI-COR Odyssey® CLx instrument (B). Relative protein levels were quantified using Fiji-Image J (85) and normalized to the untreated control. Data (mean ± range of three independent experiments) were analyzed by linear regression. Panels D,E) Effect of cobicistat on S-protein-mediated syncytia formation. Vero E6 cells were transfected with the SARS-CoV-2 S-protein and left untreated or treated with various concentrations of cobicistat or with sera isolated from convalescent SARS-CoV-2 patients (1:100 dilution). Syncytia formation was examined 24 hours post-transfection by immunofluorescence (IF) staining for DAPI and S-protein (D) and quantified as the number of cells forming syncytia (E). Data were analyzed using the nonparametric Kruskal-Wallis test followed by Dunn’s post-test. Horizontal lines represent mean values. ** P < 0.01; **** P < 0.0001. Scale bar = 50μm.

We thus proceeded to analyze the possible impact of cobicistat on other key viral proteins. To reduce the bias of the analysis, while retaining a representative model of the infection, we performed western blot analysis of Vero E6 cell lysates using previously validated patient sera to detect viral proteins (49). The results showed the reduction of a high molecular weight band (≈250 kDa) when infected cells were incubated with low micromolar concentrations of cobicistat (Supplementary Figure 4A). Based on the known molecular weights of SARS-CoV-2 proteins, we postulated that the patterns detected with patient sera corresponded to dimers/trimers of the S-protein (50,51) and to the nucleoprotein (N-protein) (51) of the virus. To confirm this hypothesis, we performed western blot analysis using monoclonal antibodies directed against the S- and N-protein (Figure 3B). The results confirmed the observation that S-protein levels are decreased by cobicistat (Figure 3B). Moreover, quantification of viral proteins in our western blots indicated a higher inhibition by cobicistat of the cleaved form (≈100 KDa) of the S-protein (50,52), which is responsible for the fusion to the host cell and subsequent viral entry (13) (Figure 3C). To isolate the possible effect of cobicistat on S-protein-mediated fusion, we used a cellular assay measuring syncytia formation in Vero E6 cells transfected with the S-protein. The results showed decreased syncytia formation when cells were incubated with cobicistat or when sera from SARS-CoV-2 patients were used as controls to block S-protein fusion (Figure 3D,E; Supplementary Figure 4B). Of note, both western blot analysis and the syncytia assay (Figure 3C-E; Supplementary Figure 4B) indicated an effect of cobicistat in the 5-10μM range, which corresponds to the range of most IC50 values calculated on the basis of viral RNA levels in supernatants (Figure 2A,C).

Overall, these data show that the antiviral effect of cobicistat is not mediated by inhibition of 3CL_pro_ activity, but is rather exerted, at least partially, through impairment of S-protein-mediated fusion.

### Cobicistat potently enhances the antiviral effect of remdesivir in cell lines and a primary colon organoid

We then tested the potential of cobicistat to exert a double activity as direct inhibitor of SARS-CoV-2 replication and as pharmacoenhancer of other antivirals. To this aim, we evaluated remdesivir as a candidate compound to synergize with cobicistat. The choice of remdesivir was motivated by its known activity as inhibitor of SARS-CoV-2 RdRP (53), as well as by its postulated susceptibility to extensive first pass liver metabolism, potentially mediated by the cellular targets of cobicistat CYP3A and P-gp [E.M.A., Human Medicines Division. Summary on compassionate use Remdesivir Gilead (35)]. We thus examined the *in-silico* predicted affinity of remdesivir for the main members of the CYP3A family (CYP3A4 and 5), as well as for P-gp. Multiple machine learning models predicted remdesivir as a potential CYP3A4 substrate (Supplementary Table 4). Moreover, the SwissADME server (54) predicted remdesivir to be both a CYP3A4 and P-gp substrate with 79% and 88% accuracy, respectively. Similarly, the pkCSM (55) and CYPreact (56) servers also predicted remdesivir to be a substrate, but not an inhibitor, of both P-gp and CYP3A4. Finally, remdesivir displayed high docking scores to the active sites of CYP3A4, CYP3A5 and P-gp, which were comparable to those of ritonavir and cobicistat, *i*.*e*. known inhibitors with well characterized binding (Supplementary Figure 5).

To identify the most suitable *in-vitro* model for testing the combination of remdesivir and cobicistat, we first examined the relative expression levels of CYP3A4, CYP3A5 and P-gp in different human tissues and cell lines susceptible to SARS-CoV-2 infection (Supplementary Figure 6, Figure 4). Both transcriptomic and qPCR analysis highlighted liver, gut and kidney as major compartments of CYP3A4/5 and P-gp expression (Supplementary Figure 6A, Figure 4A), as previously described (39,57). On the other hand, primary lung tissues were characterized by lower CYP3A4/5 and P-gp expression, while the cell line Calu-3 showed intermediate characteristics, with low CYP3A4 and high P-gp expression (Supplementary Figure 6B), in line with upregulation of the latter marker in cancer cells (58). Of note, SARS-CoV-2 infection was associated with altered expression of these genes. In this regard, cell lines of gut origin and Vero E6 cells displayed a peculiar trend showing opposite expression patterns of CYP3A4 and CYP3A5 upon infection (Figure 4B). Given their divergent response to the infection, we decided to use both Vero E6 and T84 cells as models for testing cobicistat and remdesivir, to obtain data on the efficacy of the drug combination and on its possible reliance on increased expression of either CYP3A4 or CYP3A5. While treatment with remdesivir-only displayed antiviral activity at previously described levels (Supplementary Figure 7) (6), the combined use of cobicistat and remdesivir was able to significantly enhance the effect of each drug alone, in both cell lines (Figure 5 A-F, Supplementary Figure 8A-D). In particular, the drug combination was synergistic in almost completely abrogating viral infection/replication, as measured by IF (Figure 5 A,B; Supplementary Figure 8B), and qPCR (Figure 5C-E; Supplementary Figure 8C), analysis. In line with this potent antiviral activity, the cobicistat/remdesivir combination also displayed a synergistic effect in inhibiting the cytopathic effects of SARS-CoV-2, thus restoring viability of infected cells to levels comparable to mock-infected controls (Figure 5F; Supplementary Figure 8A,D). Finally, we tested the effect of the drug combination on a primary human colon organoid (Figure 5G), which is susceptible to SARS-CoV-2 infection, as previously described (59). Also in this case, the addition of cobicistat enhanced the antiviral effect of remdesivir (Figure 5G).

**Figure 4.**
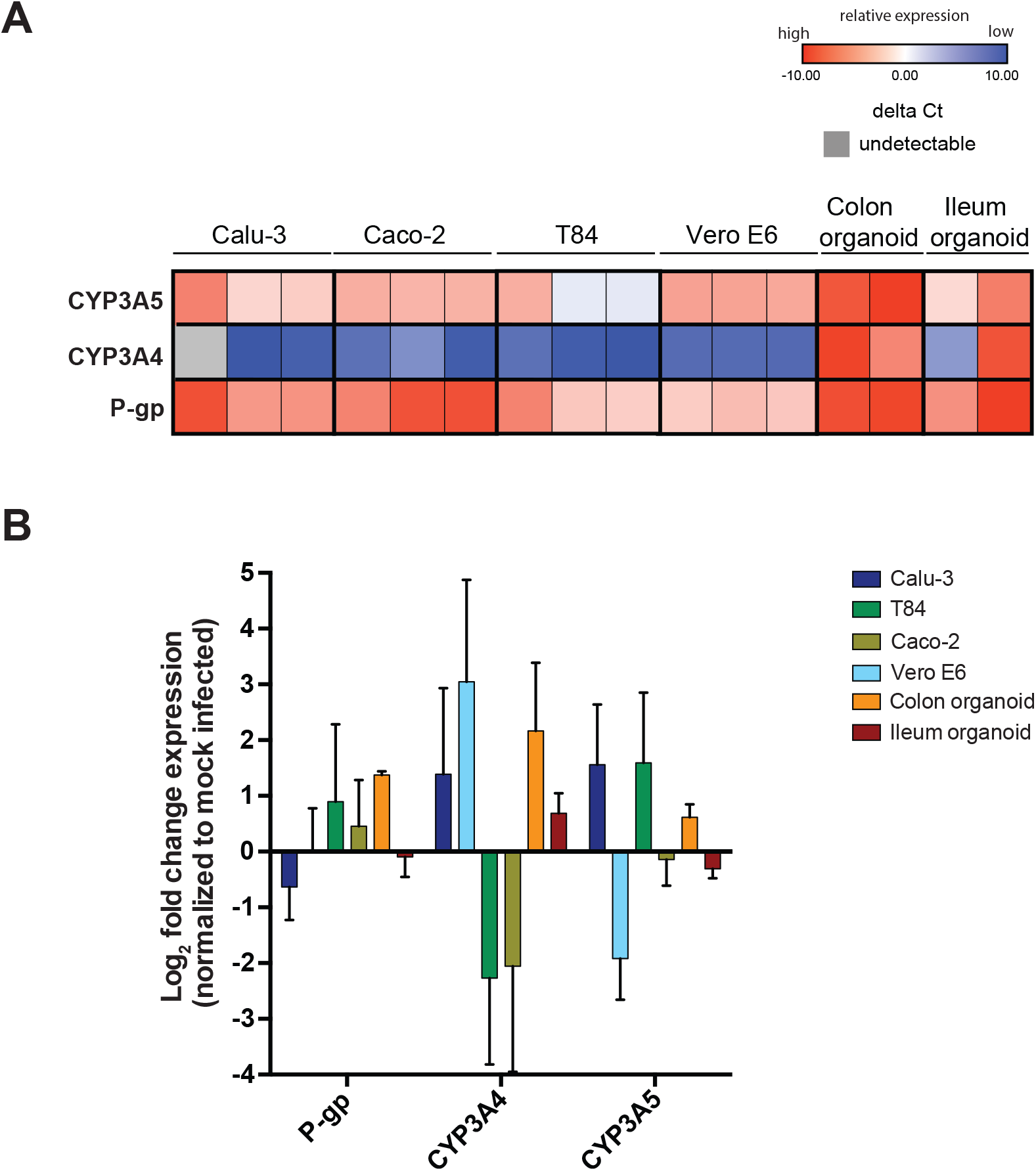
Expression of the metabolic targets of cobicistat in uninfected and SARS-CoV-2 infected cell lines and primary human organoids. Panels A,B) The relative expression of CYP3A4/5 and P-gp was analyzed by qPCR in uninfected (A) and SARS-CoV-2 infected or mock infected (B) cells. Infections were carried out at MOI 0.5 for 48 hours. Raw data were used to calculate delta CT values (A), by using the *TBP* gene as housekeeping control. Fold changes, in infected over mock infected cells (B), were calculated using the delta-delta CT method, as described in (82). Data in (B) are expressed as mean ± SD (N = 3).

**Figure 5.**
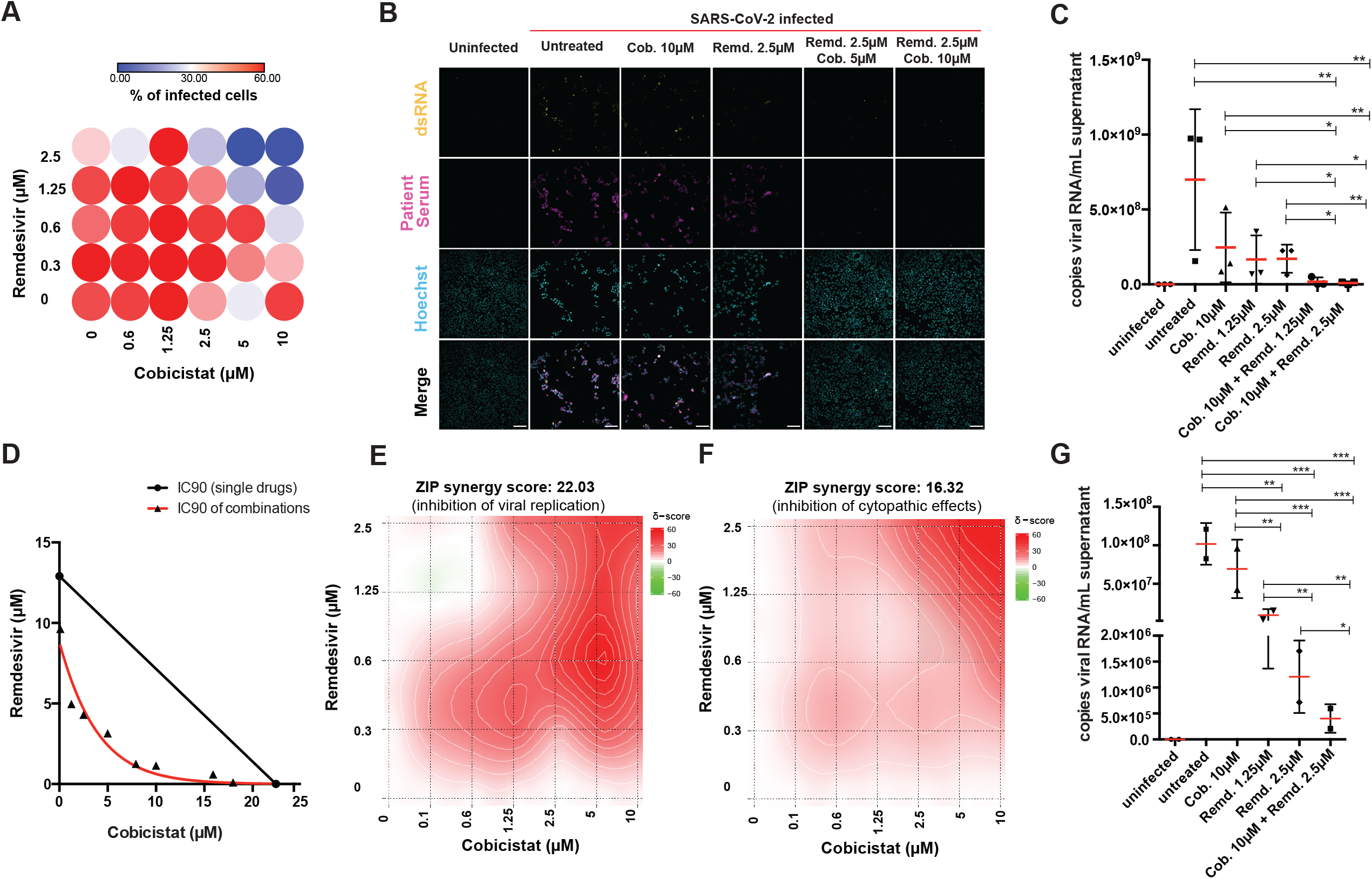
The combination of cobicistat and remdesivir synergistically inhibits SARS-CoV-2 activity. Panels A-F) Synergistic activity of cobicistat and remdesivir in inhibiting replication and cytopathic effects of SARS-CoV-2 in Vero E6 cells. Cells were infected at 0.5 MOI and left untreated or treated with the drugs at the indicated concentrations two hours-post infection. Forty-eight hours post-treatment: cells were fixed for immunofluorescence (IF) staining (A,B), supernatants were collected for qPCR (C-E) or cellular viability was analyzed (F). For IF detection, cells were stained with sera of SARS-CoV-2 patients and with the J2 antibody, which binds to double stranded RNA (49). The percentage of infected cells was determined by automatic acquisition of nine images per well (A), as described in the Methods section. Scale bar = 100μm. Viral RNA in supernatants was detected by qPCR using an *in-vitro* transcribed standard curve for absolute quantification (C-E). Data, expressed as mean ± SD, were transformed as Log_10_ to restore normality and analyzed by one-way ANOVA, followed by Holm-Sidak’s post-test (C). Cellular viability was measured by MTT assay (F). Isobologram analysis of synergism (D) (90) was performed using the IC90 values for SARS-CoV-2 replication of cobicistat, remdesivir, or their combination, calculated by non-linear regression. Synergism analyses of the inhibition of viral replication (E) or cytopathic effects (F) were performed with the SynergyFinder web-tool (86) using the Zero Interaction Potency (ZIP) model based on inhibition values calculated as described in the Methods section. Panel G) Effect of the combination of cobicistat and remdesivir on SARS-CoV-2 RNA expression in supernatants of a primary human colon organoid. Treatment with cobicistat/remdesivir was performed two hours post-infection and supernatants were collected forty-eight hours post-treatment. Viral RNA was quantified as described for panel (C). For all panels N = 3 independent experiments, except for panel E (N = 2 independent experiments) and panel G (N = 2 replicates from one colon organoid donor). *** P < 0.001; ** P < 0.01; * P < 0.05.

Overall, our data prove that the combination of cobicistat and remdesivir can suppress viral replication in different cellular models of SARS-CoV-2 infection and suggest that cobicistat can exert a double activity as direct antiviral and pharmacoenhancer.

## Discussion

The data herein presented demonstrate the antiviral activity of the FDA-approved drug cobicistat and support its possible role as a basis for combined antiviral therapies against SARS-CoV-2. The use of drug combinations targeting different steps of the viral life cycle is a well-established paradigm for treating RNA-virus infections (60,61). Translating this concept to SARS-CoV-2 drug development has, however, proven challenging due to the paucity of effective drug candidates available. In particular, compounds showing promise in initial studies, have failed to reproducibly decrease the mortality and morbidity of the infection (6–9). Similarly disappointing results were observed in the early stages of HIV-1 drug discovery, and might be partially explained by the inability of candidate antivirals to reach *in-vivo* concentrations sufficient to completely block viral replication. The use of pharmacoenhancers such as cobicistat (40) could help to overcome this limitation. While the present study exclusively focused on the combination of cobicistat and remdesivir, more than 30% of all drugs are metabolized by the main cellular targets of cobicistat (*i*.*e*. CYP3A4/5)(62). For example, the recently described SARS-CoV-2 inhibitor plitidepsin (63) is mainly metabolized by CYP3A4 *in vitro (64)*. Therefore, it is conceivable that a synergistic effect similar to that described for remdesivir might be obtained by coupling cobicistat to other antiviral agents. In particular, other compounds tested in clinical trials of SARS-CoV-2 patients, such as chloroquine/hydroxychloroquine (65) and lopinavir (62) are well known substrates of CYP3A. The booster effect of cobicistat would be further complemented by the own antiviral activity of this drug, which was here proven on several models of SARS-CoV-2 infection. In line with this, we observed the strongest synergistic effect with remdesivir, when cobicistat was used at concentrations above its IC50 levels, suggesting a combination of pharmacokinetic and pharmacodynamic effects. Of note, the concentration range in which cobicistat could inhibit SARS-CoV-2 replication was higher than that achievable through standard dosages (*i*.*e*. 150mg/day) approved for treatment of HIV-1 infection (37). This potential drawback could be mitigated by the fact that cobicistat (as well as its “parent” drug, ritonavir) was previously shown to be well tolerated at much higher concentrations, both in mice and humans. Indeed, plasma levels achievable with a single administration of high-dose (*i*.*e*. 400mg/day) (46) cobicistat in humans are predicted to overlap with concentrations displaying antiviral activity *in vitro*. These observations can also explain the limited effects or outright lack of success of early trials testing the HIV-1 protease inhibitor darunavir, boosted by a standard cobicistat dose (20,21). It is important to note that drug regimens for HIV-1 treatment are based on the assumption of long-term (or even lifelong) administration, while antiviral treatment for SARS-CoV-2 typically spans between one and two weeks. Thus, an increase of the cobicistat dosage in the context of an acute illness such as COVID-19 could be considered, because its administration would be transient.

Another possible limitation of candidate antivirals for SARS-CoV-2 treatment is the inability to reach specific tissue reservoirs of the infection. Remdesivir is case in point, due to its quick metabolization and poor intestinal absorption (66). Of note, previous experience with HIV-1 protease inhibitors suggests that cobicistat might overcome this limitation (67), in line with the synergistic effect that we observed when treating primary colon organoid and T84 colon adenocarcinoma cells with the combination of cobicistat and remdesivir. Intriguingly, the tissue penetration and activity of cobicistat in the main sites of CYP3A expression (*i*.*e*. gut and liver) might be relevant also for the route of administration of remdesivir. Currently, remdesivir requires intravenous administration due to its extensive first pass metabolism (68), but it is conceivable that coupling it with cobicistat might improve its absorption, perhaps allowing oral formulation of the drug. Increasing the scalability of remdesivir might *per se* improve its therapeutic potential, as an early treatment of the infection might prevent hospitalization and development of severe COVID-19, a stage where the efficacy of remdesivir could not be firmly established(8).

An important advantage of antiviral combinations is the possibility to target multiple steps of the viral life cycle. Despite the *in-silico* docking results of our study and of other groups (43–45), our *in-vitro* data demonstrate that cobicistat does not inhibit the enzymatic activity of 3CL_pro_. Our molecular dynamics analysis suggests that this discrepancy might be reconciled when conformational entropy (-TΔS) is included among the analysis parameters, which reduces the expected binding energy between cobicistat and 3CL_pro_. On the other hand, our *in-vitro* results suggest an effect of cobicistat on S-protein maturation or function. While the results do not allow distinguishing between these two potential effects, the time course analysis suggests that the antiviral activity of cobicistat might be exerted following the first cycle of SARS-CoV-2 cellular entry. Other potential explanations of our results could be an effect of cobicistat in inhibiting the cleavage of the S-protein by furin (69) or by other proteases, or in affecting the turnover or expression of ACE-2. Further studies will be required to precisely characterize the molecular mechanism of antiviral action of cobicistat.

One limitation of our work is that it does not specifically examine the metabolization of remdesivir in the presence or absence of cobicistat. Of note, hydrolases are expected (more than CYP3A or P-gp) to play a paramount role in the metabolization of remdesivir *in vivo (70)*. On the other hand, published evidence on the metabolic conversion of remdesivir is scarce, and most metabolic considerations rely on generic guidance about expected drug-drug interactions (70). Another aspect that could limit the applicability of our results is the possibility that cobicistat may not reach effective concentrations with clinically acceptable doses *in vivo*. While only animal or clinical studies will be able to test this question, previous results on the tissue distribution and plasma concentrations of cobicistat in mice and humans suggest that low-micromolar concentrations of cobicistat might reach the main sites of SARS-CoV-2 replication using well-tolerated dosages (Pharmacology Review of Cobicistat - Application number: 203-094) (46,71).

Overall, our study introduces cobicistat as a new candidate for inhibiting SARS-CoV-2 replication and for designing combination therapies aimed at blocking or reversing the onset of COVID-19.

## Materials and Methods

### Virtual screening and molecular docking

Identification of potentially active SARS-CoV-2 inhibitors with desirable Absorption, Distribution, Metabolism, Excretion and Toxicity (ADME-Tox) properties, was performed by structure-based virtual screening (SBVS) of Drugbank V. 5.1.5(72) compounds targeting the three-dimensional structure of SARS-CoV-2 3CL_pro_. The analysis was focused on the substrate-binding site, which is located between domain I and II of 3CL_pro_. The binding site was identified using the publicly available 3D crystal structure [Protein Data Bank (PDB) ID: 6W63]. Structures of the previously described non-covalent protease inhibitor X77(73), natively co-crystallized with 3CL_pro_ were used as a reference for the identification of binding-site coordinates and dimensions for the virtual screening workflow, as well as for the docking validation of positions generated from the screening.

Protein structure analysis and preparation for docking were performed using the Schrödinger protein preparation wizard (Schrödinger Inc). Missing hydrogen atoms were added, bond orders were corrected and unknown atom types were assigned. Protein side-chain amides were fixed using program default parameters and missing protein side chains were filled-in using the prime tool. All non-amino acid residues, including water molecules, were removed. Further, unrelated ligand molecules were removed and active ligand structures were extracted and isolated in separate files. Finally, the minimization of protein strain energy was achieved through restrained minimization options with default parameters. The centroids of extracted ligands were then used to identify the binding site with coordinates and dimensions extended for 20 Å stored as Glide grid file. Drug screening was performed using the Glide software (74). High throughput virtual screening (HTVS) was performed with the fastest search configurations. After post-docking minimization, the top-scoring tenth percentile of the output docked structures were subjected to the standard precision docking stage (SP). Then, active ligand structures were extracted and isolated in separate files. Finally, the top 10% scoring compounds were selected and retained only if their good scoring states were confirmed by Extra precision docking.

Remdesivir docking to CYP3A4, CYP3A5 and P-gp structures was performed to assess its capacity as a substrate/inhibitor for these proteins. CYP3A4, CYP3A5 and P-gp structures were collected from Protein Data Bank (PDB), IDs: 5VC0, 5VEU and 6QEE, respectively, and were subjected to the same preparation steps described in the former section. Native inhibitors were used for identification of binding sites; the centroid of the known inhibitor Zosuquidar was used to identify the drug binding pocket of the P-gp protein structure. Further, co-crystallized Ritonavir was used for identification of the drug binding pocket in both CYP3A4/5. Receptor grids were generated for protein structures, for both CYP3A4 and CYP3A5. The heme iron of the Protoporphyrin ring was added as metal coordination constraint, allowing metal-ligand interaction in the subsequent docking steps. Docking was performed using flexible ligand conformer sampling allowing ring sampling with a 2.5 kcal/mol window. Retained poses for the initial docking phase were set to 5000 poses and only 800 best poses per ligand were selected for energy minimization. Finally, post-docking minimization was carried out for 10 poses per ligand with a 0.5 kcal/mol threshold for rejecting minimized poses.

### Molecular dynamics

Candidate ligand-receptor complexes derived from docking simulations were chosen for further computational analysis through molecular dynamics. The complexes examined included SARS-CoV-2 3CL_pro_ protein bound to cobicistat, darunavir, X77, nelfinavir, ritonavir, tipranavir, GC376, lopinavir, and MG-132. The complex with the native co-crystallized binding ligand, X77, obtained from the PDB structure with the code 6W63 was used for comparison in addition to the docked X77. The ligand receptor complexes were protonated and processed via Molecular Operating Environment (MOE 2012; Chemical Computing Group). The AMBER 18 molecular dynamics package was used for the molecular dynamics simulations. The force field AMBER ff14SB (75) was used for the protein while the force field GAFF2 (76) was used for the ligands. Each complex was solvated in a cubic box extending 15 Å in each direction. The system was neutralized by the addition of Na^+^ ions followed by the addition of extra Na^+^ and Cl^−^ to bring the salt concentration to 150 mM. The system was energy minimized using a series of steepest descent and conjugate gradient minimization steps followed by a series of constant pressure equilibration runs under decreasing position restraints, from 5.0 to 0.1 kcal mol^-1^Å^-2^. After that, an unrestrained production run of 30 ns was performed. All the dynamics were performed in the NPT ensemble at 310K employing Langevin thermostat and Berendsen barostat with a non-bonded cutoff value of 8.0 Å. The trajectories were saved every 10 ps and analyzed for RMSD equilibration using CPPTRAJ (77). After that, the binding energy between the ligand and the receptor was estimated using Molecular Mechanics / Generalized-Born Surface Area (MM/GBSA) method as implemented in AmberTools (78). 300 snapshot/ns were used for the binding energy estimation. The IGB model 8 was used at a salt concentration of 100 mM. Other parameters were left at their default values. The entropic contribution was estimated using a previously described method of calculation of interaction entropy (48,79).

### Cell lines and primary human organoids

The following cell lines were used for infection and/or relative quantification of gene expression: Calu-3 (ATCC HTB-55), Caco-2 (ATCC HTB-37), T84 (ATCC CCL-248) and VeroE6 (ATCC CRL-1586). Primary organoids derived from human colon and ileum were seeded in 2D as described in (59). Culture conditions and susceptibility to SARS-CoV-2 infection have been previously described (59,80).

### Virus stock production and infection

Viral stocks used for infections were produced by passaging the BavPat1/2020 SARS-CoV-2 strain in Vero E6 cells and the infectious titer was estimated by plaque assay, as previously described (49). Infection experiments were conducted using 2.5 × 10^4^ or 2.5 × 10^5^ cells per well in 96 and 12 well plates, respectively. Cell lines were infected at 0.05 or 0.5 MOI in medium with low FCS content (2%). Colon organoids were infected in a 24-well plate using 6 × 10^4^ plaque forming units (PFU) per well. Two hours post-infection cells were washed twice in PBS and resuspended in complete medium.

### Drug treatments

The following compounds were tested to determine their effects on 3CL_pro_ activity, cytotoxicity or inhibition of SARS-CoV-2 replication: cobicistat (#sc-500831; Santa Cruz Biotechnology), remdesivir (#S8932; Selleckchem Chemicals), tipranavir (#sc-220260; Santa Cruz Biotechnology), nelfinavir mesylate hydrate (#PZ0013, Sigma-Aldrich), darunavir, lopinavir (both obtained through the AIDS Research and Reference Reagent Program, Division of AIDS, NIAID), MG-132 (#M8699; Sigma-Aldrich), GC376 (BPS Bioscience).

### RNA isolation and cDNA retrotranscription

RNA extraction was performed on cell lysates or supernatants using the NucleoSpin RNA, Mini kit for RNA purification (Macherey-Nagel, Düren, Germany) according to the manufacturer’s instructions. The concentration of RNA extracted from cell lysates was measured using a P-class P 300 NanoPhotometer (Implen GmbH, Munich, Germany). Retrotranscription to cDNA was performed with 500ng of intracellular RNA or 10μL of RNA from supernatants, using the High-Capacity cDNA Reverse Transcription Kit (Applied Biosystems, Foster City, CA, USA) following the manufacturer’s instructions.

### SARS-CoV-2 RNA standard

For the preparation of a viral RNA standard to use in qPCR for quantification of viral copies in supernatants, SARS-CoV-2 N sequence was reverse transcribed from total RNA isolated from cells infected with the SARS-CoV-2 BavPat1 stain using Superscript 3 and specific primers (TTAGGCCTGAGTTGAGTCA). The resulting cDNA was amplified and cloned into the pJET1.2 plasmid. Plasmid DNA (10 μg) was linearized by AdeI restriction enzyme digestion and DNA was purified using the NucleoSpin Gel and PCR Clean-up kit (Macherey-Nagel, Düren, Germany). For *in-vitro* transcription, T7 RNA polymerase was used as previously described(81). *In-vitro* transcripts were purified by phenol-chloroform extraction and resuspended in RNase-free water. RNA integrity was confirmed by agarose gel electrophoresis.

### qPCR analysis

Gene and/or viral expression were analyzed by SYBR green qPCR using, for each reaction, 10μL of SsoFast™ EvaGreen® Supermix (Bio-Rad Laboratories, Hercules, CA, USA), 500nM of forward and reverse primer (0.1μL each from 100μM stock), 8.8μL water and 1μL cDNA. The primers used are listed in Supplementary table 2. The qPCR reaction was performed on a CFX96/C1000 Touch qPCR system (Bio-Rad Laboratories, Hercules, CA, USA) using the following PCR program: polymerase activation/DNA denaturation 98°C for 3 min, followed by 45 cycles of denaturation at 98°C for 10s; annealing/extension at 60°C for 40s and a final extension step at the end of the program at 65°C for 30s. Gene expression data were normalized using the 2(−ΔΔ C(T)) method (82), using the Tata-binding protein (*TBP*) gene as housekeeper control.

### Western blot

For Western blot experiments 0.5 × 10^6^ cells were lysed in a buffer (20 mM Tris–HCl, pH 7.4, 1 mM EDTA, 150 mM NaCl, 0.5% Nonidet P-40, 0.1% SDS, and 0.5% sodium deoxycholate) supplemented with protease and phosphatase inhibitors (Sigma-Aldrich, Saint Louis, MI, USA). Lysates were supplemented with loading buffer, boiled at 95°C for 10 min and sonicated for 5 min using a Bioruptor® Plus sonication device (Diagenode, Liège, Belgium). Protein lysates were then run on a precast NuPAGE Bis-Tris 4–12% SDS–PAGE (Thermo Fisher Scientific, Waltham, MA, USA) and transferred onto a nitrocellulose membrane (GE Healthcare, Little Chalfont, UK) using a Trans–Blot device for semi-dry transfer (Bio-Rad Laboratories, Hercules, CA, USA). Membranes were blocked using the LI-COR Intercept (PBS) Blocking Buffer (LI-COR Biosciences, Lincoln, NE, USA) for 1 h at RT and incubated overnight at 4°C with the following primary antibodies diluted in blocking buffer + 0.2% Tween 20: α-β-actin (1:10,000), (Sigma-Aldrich, Saint Louis, MI, USA), α-SARS-CoV-2 S-protein [(rabbit; 1:1000) ab252690 Abcam], α-SARS-CoV-2 N-protein [(mouse; 1:1000) AB_2827977, Sino Biological)], sera of SARS-CoV-2 convalescent individuals (1:200). Sera were collected as described in(49), following signing of informed consent by the donors, as well as ethical approval by Heidelberg University Hospital. After primary antibody incubation, membranes were washed three times with 0.1% PBS-Tween and incubated for 1 h with the following fluorescence-conjugated secondary antibodies: IRDye® 800CW Goat anti-Human IgG, IRDye® 800CW anti rabbit, IRDye® 700CW anti mouse (LI-COR Biosciences, Lincoln, NE, USA). All secondary antibodies were diluted 1:15000 in blocking buffer + 0.2% Tween. After three washes with 0.1% PBS-Tween and one wash in PBS, fluorescence signals were acquired using a LI-COR Odyssey® CLx instrument.

### Re-processing of microarray and RNA-Seq data

Microarray gene expression data for CYP3A4/5 and P-gp in different anatomical tissues or cell lines were retrieved from the Homo Sapiens Affymetrix Human Genome U133 Plus 2.0 Array dataset. Data were filtered by applying the criteria “Healthy sample status” and “No experimental treatment”. From the initial list, tissues with sample size < 25 were filtered out. The anatomy search tool was used to plot Log_2_ expression ratios of the tested genes. Gene expression data in cell lines were retrieved from the aforementioned microarray dataset and from the RNAseq “mRNA Gene Level Homo sapiens (ref: Ensembl 75)” dataset. The cell line condition filter was used to refine the analysis and include exclusively cell lines susceptible to SARS-CoV-2 infection (*i*.*e*. T84, Caco2, Calu-3 and A-549).

### Cell viability

Cell viability was evaluated by (3-[4,5-dimethylthiazol-2-yl]-2,5 diphenyl tetrazolium bromide) (MTT) assay and by crystal violet staining as previously described(83,84). Briefly, the MTT assay was conducted using the CellTiter 96® Non-Radioactive Cell Proliferation Assay (MTT) (Promega; Madison, WI, USA). Cells were plated in a 96-well plate at a concentration of 3 × 10^6^ cells/mL in 100 μl of medium. The MTT solution (15 μl) was added to each well and, after 2–4 h, the reaction was stopped by the addition of 100 μl of 10% SDS. Absorbance values were acquired using an Infinite 200 PRO (Tecan, Männedorf, Switzerland) multimode plate reader at 570 nm wavelength. For crystal violet staining, cells were fixed in 6% formaldehyde and incubated with 0.1% crystal violet for 15 mins. Unbound staining was then washed with H_2_O and cells were imaged using a Nikon Eclipse Ts2-FL microscope.

### 3CL_pro_ Fluorescence Resonance Energy Transfer (FRET) assay

The activity of 3CL_pro_ was measured by FRET assay (BPS Bioscience, San Diego, CA, USA) according to the manufacturer’s instructions and as previously described (47). Briefly, serial dilutions of test compounds and known 3CL_pro_ inhibitors were incubated in a 384 well plate with the 3CL_pro_ enzyme and its appropriate buffer, containing 0.5 M DTT. Wells without drugs or without 3CL_pro_ were used as positive control of 3CL_pro_ activity and blank control, respectively. After a 30 min incubation, the 3CL_pro_ substrate was added to each well and the plate was stored for 4 hours in the dark. The fluorescence signal was acquired on an Infinite 200 PRO (Tecan, Männedorf, Switzerland) using an excitation wavelength of 360nm and a detection wavelength of 460nm. Relative 3CL_pro_ was expressed as percentage of the positive control after subtraction of the blank.

### Immunofluorescence

For immunofluorescent (IF) staining, cells were seeded on iBIDI glass bottom 96 well plates and infected with SARS-CoV-2 the BavPat1/2020 strain at MOI 0.5. Cells were rinsed in PBS and fixed with 6% formaldehyde, followed by permeabilization with 0.5% Triton X100 (Sigma) in PBS for 15 min. Cells were then blocked in 2% milk (Roth) in PBS and incubated with primary antibodies in PBS (anti dsRNA mouse monoclonal J2 antibody (Scicons) 1:2000 and convalescent SARS-CoV-2 patient serum 1:250). Afterwards, cells were washed twice in PBS 0.02% Tween and incubated with secondary antibodies in PBS [1:1000 anti-mouse 568, Goat anti-human IgG-AlexaFluor 488 (Invitrogen, Thermofisher Scientific) for detection of human immunoglobulins in serum and goat anti-mouse IgG-AlexaFluor 568 (Invitrogen, Thermofisher Scientific) for dsRNA detection]. Nuclei were counterstained with Hoechst 33342 (Thermofisher Scientific, 0.002µg/ml in PBS) for 5 min, washed twice with PBS and stored at +4°C until imaging.

### Microscopy and image analysis

Cells were imaged using a motorized Nikon Ti2 widefield microscope or with a Nikon/Andor (CSU W1) spinning disc using a Plan Apo lambda 20x/0.75 air objective and a back-illuminated EM-CCD camera (Andor iXon DU-888). The JOBS module was used for automatic acquisition of 9 images per well. Images were acquired in 3 channels using the following excitation/emission settings: Ex 377/50, Em 447/60 (Hoechst); Ex 482/35, Em 536/40 (AlexaFluor 488); Ex 562/40, Em 624/40 (AlexaFluor 568). When the spinning disc was used the excitation was performed with 405nm, 488nm and 561nm lasers.

Quantification of infected cells (expressed as percentage of total cells imaged per well) was performed using a custom-made macro in ImageJ (85). After camera offset subtraction and local background subtraction using the rolling ball algorithm, nuclei were segmented using automated local thresholding based on the Niblack method. Region of interest [represented by the ring (5 pixel wide) around the nucleus] was determined for each individual cell. Median signal intensity was measured in the region of interest in Alexa488 (convalescent SARS-CoV-2 serum) and Alexa568 (dsRNA) channels. Threshold for calling infected cells was manually determined for each individual experiment using the data from mock infected cells. The same image analysis procedure and threshold was used for all wells within one experiment.

### Syncytia formation assay

For the syncytia formation assay, Vero E6 cells (0.2 × 10^6^ cells/well) were seeded on cover slips in a 12 well plate 24 h before transfection. Cells were transfected using TransIT-2020 or TransIT-LT1 (Mirus) with 0.75 µg of pCDNA3.1(+)-SARS-CoV-2-S and 100 µl Opti-MEM per well. 2 h post transfection, cells were treated with cobicistat (final concentration of 1 µM, 5 µM and 10 µM), sera of convalescent SARS-CoV-2 patients (1:500 or 1:100) or DMSO. 24 h post transfection, cells were washed twice with PBS and fixed in 4 % paraformaldehyde (PFA) for 20 min at RT. After another washing step, cells were permeabilized in 0.5 % Triton for 5 min at RT, washed and blocked in 3% lipid-free BSA in PBS-0.1% Tween-20 for 1 h at RT. After washing, cells were stained with the primary rabbit polyclonal anti-SARS-CoV-2 spike glycoprotein antibody (1:1000, Abcam) for 1 h at RT or overnight at 4 °C. After washing, cells were incubated with the secondary Alexa Fluor 488 goat anti-rabbit IgG antibody (1:500, Life Technologies) for 1 h at RT. Cells were then washed again and incubated with DAPI (1:1000, Sigma-Aldrich) for 1 min followed by washing with PBS and deionized H_2_O. Images were acquired with a Nikon Eclipse Ts2-FL Inverted Microscope. Syncytia with three or more nuclei surrounded by the antibody staining were considered for the quantification. The edges of the antibody staining were overdrawn with the polygon selection tool in Image J (85).

### Statistical analysis

Data normality assumptions were tested by D’Agostino & Pearson normality test (for N > 3). Multiple group comparisons were conducted by non-parametric Kruskal-Wallis test, followed by Dunn’s post-test, or by one-way or two-way ANOVA followed by Holm-Sidak’s or Dunnett’s post-tests, respectively. Half maximal inhibitory (IC50), effective (EC50) and cytotoxic (CC50) concentrations of the compounds were estimated by nonlinear regression after data normalization. For synergy and for IC50 calculation the normalized relative inhibition values were calculated according to the formula: % inhibition = 100 * (1 - (X - mock infected)/(infected untreated - mock infected)), where X is each given treatment condition. Data analysis was conducted using GraphPad Prism v6 (GraphPad Software, San Diego, CA, USA). Synergy scores were calculated with the SynergyFinder web-tool(86) using the Zero Interaction Potency (ZIP) model (87).

## Authors’ Contributions

I.L.S., M.M.T., M.F., A.S. conceived the project. I.L.S, A.S., M.L. directed the project. I.L.S., M.C., C.N., P.C., M.S., O.T.F., S.B., R.B., A.S., M.L. designed the experiments. M.M.T., M.F., A.A. performed *in-silico* analyses. I.L.S, B.L., L.G., L.Z., M.S., A.S., M.L. performed *in-vitro* experiments. B.L., L.Z., V.L., M.L. performed microscopy analysis. I.L.S., B.L., L.G., L.Z. analyzed *in-vitro* data. I.L.S. wrote the manuscript.

## Acknowledgments

The authors thank Dr. Stefan Pöhlmann (DPZ Göttingen) for providing the pCDNA3.1(+)-SARS-CoV-2-S plasmid. I.L.S. is supported by the Fundação de Amparo à Pesquisa do Estado de São Paulo (CNPJ 43.828.151/0001-45). O.T.F acknowledges support from the Deutsche Forschungsgemeinschaft (Projektnummer 240245660 – SFB 1129), MWK Baden-Württemberg (Sonderfördermaßnahme COVID-19 Forschung project HD18).

## Supplementary Figures

**Supplementary Figure 1. Validation of the antiviral potential of cobicistat**. Panels A,B) Validation of the antiviral effects of cobicistat shown in Figure 1D,E using a distinct set of primers (2019-nCoV_N2 primer set; Center of Disease Control). Fold change values in intracellular RNA (A) were calculated by the delta-delta CT method (82), using the Tata-binding protein (TBP) gene as housekeeper control. Expression levels in supernatant (B) were quantified using an *in-vitro* transcribed standard curve generated as described in the Methods section. Data are expressed as mean with SD and were analyzed by two-way ANOVA followed by Dunnet′s post-test (N = 3 independent experiments). * P < 0.05; ** P < 0.01.

**Supplementary Figure 2. Effect of various concentrations of cobicistat on the viability of the cell lines employed in the study**. Panels A-C) Uninfected cell lines of lung (Calu-3; A), gut (T84; B) and kidney (Vero E6; C) origin were left untreated or treated with serial dilutions of cobicistat. Forty-eight hours post-treatment cellular viability was measured by MTT assay. Data, expressed as mean ± SD of three independent experiments, were normalized to the untreated control and CC50 values were calculated by nonlinear regression.

**Supplementary Figure 3. *In-silico* prediction of the binding stability over time of potential 3CL**_**pro**_ **ligands**. Binding stability of the ligands to 3CL_pro_ was estimated by molecular dynamics. The previously described inhibitors of 3CLpro X77 (73), GC376 and MG-132 (89) were included as comparisons.

**Supplementary Figure 4. Validation of the impact of cobicistat treatment on S-protein expression and fusion**. Panel A) Effect of cobicistat treatment on SARS-CoV-2 protein expression in infected Vero E6 cells. Cells were infected at 0.5 MOI and left untreated or treated, two hours post-infection, with various concentrations of cobicistat, of the RdRP inhibitor remdesivir, or the 3CL_pro_ inhibitor GC376. Cells were harvested 24 hours post-treatment and subjected to protein extraction and Western Blot analysis. Expression of viral proteins was detected using sera from convalescent SARS-CoV-2 patients (at 1:200 dilution) followed by incubation with a fluorescent-conjugated anti-human secondary antibody. Expression of the housekeeping protein actin-β was detected using a specific primary monoclonal antibody. Fluorescent signals were measured using a LI-COR Odyssey® CLx instrument. Panel B) Quantification of the inhibition of S-protein-mediated syncytia formation by cobicistat. Vero E6 cells were transfected with the SARS-CoV-2 S-protein and left untreated or treated with various concentrations of cobicistat or with sera isolated from convalescent SARS-CoV-2 patients (1:100 or 1:500 dilution). Syncytia formation was examined 24 hours post-transfection by immunofluorescence (IF) staining for DAPI and S-protein (as shown in Figure 3D) and quantified as the number of cells forming syncytia. Data were analyzed using the nonparametric Kruskal-Wallis test followed by Dunn’s post-test. Horizontal lines represent mean values. *P < 0.05; ** P < 0.01; *** P < 0.001.

**Supplementary Figure 5. Predicted *in-silico* binding of remdesivir to CYP3A4/5 and P-gp**. Panels A-C) Molecular docking analysis of the binding pose and docking score of remdesivir (DB14761) to CYP3A4 [PDB ID: 5VC0; (A)], CYP3A5 [PDB ID: 5VEU; (B)] and P-gp [PDB ID: 6QEE; (C)]. Docking scores of the CYP3A and P-gp inhibitors cobicistat (DB09065) and ritonavir (DB00503) are provided as comparisons. All analyses were conducted using the Schrödinger software package.

**Supplementary Figure 6. Expression of the metabolic targets of cobicistat in human tissues and cell lines**. Panels A,B. Expression levels of CYP3A4/5 and P-gp in different human tissues from healthy donors (A) or cell lines susceptible to SARS-CoV-2 infection (B). Gene expression data were retrieved from the “Homo Sapiens Affymetrix Human Genome U133 Plus 2.0 Array” and from the RNAseq “mRNA Gene Level Homo sapiens (ref: Ensembl 75)” datasets.

**Supplementary Figure 7. Antiviral effect of remdesivir on the Vero E6 and T84 cell lines**. Panels A,B) Effect of serial dilutions of remdesivir on SARS-CoV-2 RNA amount in supernatants and on the viability of uninfected Vero E6 (A) and T84 (B) cells. Cells were left uninfected or infected with SARS-CoV-2 using a 0.5 MOI. Infected cells were left untreated or treated with remdesivir two hours post-infection. Forty-eight hours post-infection supernatants were collected and viral RNA was assayed by qPCR. Cellular viability was measured by MTT assay in uninfected cells 48 hours post-treatment with different concentrations of remdesivir. Inhibition of viral replication and cell viability were normalized to the untreated control and half maximal inhibitory (IC50) was calculated by nonlinear regression.

**Supplementary Figure 8. Synergistic antiviral effect of cobicistat and remdesivir in the Vero E6 and T84 cell lines**. Panels A-D) Effect of combined treatment of cobicistat and remdesivir on the viability of SARS-CoV-2 infected Vero E6 cells (A) and on viral replication (B,C) and inhibition of cytopathic effects (D) in T84 cells. Cells were infected at 0.5 MOI and left untreated or treated with the drugs at the indicated concentrations two hours-post infection. Forty-eight hours post-treatment: cells were fixed for crystal violet (A) or immunofluorescence (IF) (B) staining, supernatants were collected for qPCR (C), or cellular viability was analyzed (D). For IF detection, cells were stained with sera of SARS-CoV-2 patients (B). Viral RNA in supernatants was detected by qPCR using an *in-vitro* transcribed standard curve for absolute quantification. Data, expressed as mean ± SD, were transformed as Log_10_ to restore normality and analyzed by one-way ANOVA, followed by Holm-Sidak’s post-test (C). Scale bars = 100μm. Cellular viability was measured by MTT assay and synergism analysis of the inhibition of cytopathic effects was performed with the SynergyFinder web-tool (86) using the Zero Interaction Potency (ZIP) model based on inhibition values calculated as described in the Methods section. N = 3 independent experiments (C,D). *** P < 0.001; ** P < 0.01.

**Supplementary Movie 1. Animation of the molecular dynamics trajectory of cobicistat bound to the 3CL**_**pro**_ **of SARS-CoV-2**. Molecular dynamics analysis performed including the contribution of entropy, as previously described (79), shows instability of the binding of cobicistat, characterized by its constant change in orientation.

**Supplementary Table 1. Top-scoring list of FDA-approved drugs predicted to bind 3CL**_**pro**_ ***in silico***. The Drugbank library of compounds was screened by molecular docking based on the predicted binding mode and affinity of each compound to the allosteric active site of SARS-CoV-2 3CL_pro_. Docking scores were calculated using Glide (74).

**Supplementary Table 2. List of qPCR primers used in the study**.

**Supplementary Table 3. *In-silico* and *in-vitro* affinity of cobicistat and other putative inhibitors to SARS-CoV-2 3CL**_**pro**_. *In-silico* binding stability of the ligands to 3CL_pro_ was estimated by molecular dynamics including the contribution of entropy, as previously described (79). Binding free energies (ΔGb) were calculated as described in the methods section. *In-vitro* inhibition of 3CL_pro_ was measured by FRET assay, as shown in Figure 3A. Data were normalized to the untreated control and half maximal effective concentration (EC50) values for each ligand were calculated by nonlinear regression.

**Supplementary Table 4. Machine learning prediction of potential binding of remdesivir to CYP3A4**. The likelihood of remdesivir being a substrate of CYP3A4 was estimated using the algorithms described in (54,56) and their performance was compared to that of previously described algorithms (91–98) as listed in the table.

## References

1. V’kovski P, Kratzel A, Steiner S, Stalder H, Thiel V. Coronavirus biology and replication: implications for SARS-CoV-2. Nat Rev Microbiol [Internet]. 2020 Oct 28; Available from: http://dx.doi.org/10.1038/s41579-020-00468-6

2. Pushpakom S, Iorio F, Eyers PA, Escott KJ, Hopper S, Wells A, et al. Drug repurposing: progress, challenges and recommendations. Nat Rev Drug Discov. 2019 Jan;18(1):41–58.

3. Johnson RM, Vinetz JM. Dexamethasone in the management of covid -19 [Internet]. BMJ. 2020. p. m2648. Available from: http://dx.doi.org/10.1136/bmj.m2648

4. RECOVERY Collaborative Group, Horby P, Lim WS, Emberson JR, Mafham M, Bell JL, et al. Dexamethasone in Hospitalized Patients with Covid-19 - Preliminary Report. N Engl J Med [Internet]. 2020 Jul 17; Available from: http://dx.doi.org/10.1056/NEJMoa2021436

5. Titanji BK, Farley MM, Mehta A, Connor-Schuler R, Moanna A, Cribbs SK, et al. Use of Baricitinib in Patients with Moderate and Severe COVID-19. Clin Infect Dis [Internet]. 2020 Jun 29; Available from: http://dx.doi.org/10.1093/cid/ciaa879

6. Wang M, Cao R, Zhang L, Yang X, Liu J, Xu M, et al. Remdesivir and chloroquine effectively inhibit the recently emerged novel coronavirus (2019-nCoV) in vitro. Cell Res. 2020 Mar;30(3):269–71.

7. Beigel JH, Tomashek KM, Dodd LE, Mehta AK, Zingman BS, Kalil AC, et al. Remdesivir for the Treatment of Covid-19 - Final Report. N Engl J Med. 2020 Nov 5;383(19):1813–26.

8. Wang Y, Zhang D, Du G, Du R, Zhao J, Jin Y, et al. Remdesivir in adults with severe COVID-19: a randomised, double-blind, placebo-controlled, multicentre trial. Lancet. 2020 May 16;395(10236):1569–78.

9. RECOVERY Collaborative Group, Horby P, Mafham M, Linsell L, Bell JL, Staplin N, et al. Effect of Hydroxychloroquine in Hospitalized Patients with Covid-19. N Engl J Med. 2020 Nov 19;383(21):2030–40.

10. Pawlotsky J-M, Feld JJ, Zeuzem S, Hoofnagle JH. From non-A, non-B hepatitis to hepatitis C virus cure. J Hepatol. 2015 Apr;62(1 Suppl):S87–99.

11. Gulick RM, Flexner C. Long-Acting HIV Drugs for Treatment and Prevention [Internet]. Vol. 70, Annual Review of Medicine. 2019. p. 137–50. Available from: http://dx.doi.org/10.1146/annurev-med-041217-013717

12. Guy RK, Kiplin Guy R, DiPaola RS, Romanelli F, Dutch RE. Rapid repurposing of drugs for COVID-19 [Internet]. Vol. 368, Science. 2020. p. 829–30. Available from: http://dx.doi.org/10.1126/science.abb9332

13. Hoffmann M, Kleine-Weber H, Schroeder S, Krüger N, Herrler T, Erichsen S, et al. SARS-CoV-2 Cell Entry Depends on ACE2 and TMPRSS2 and Is Blocked by a Clinically Proven Protease Inhibitor. Cell. 2020 Apr 16;181(2):271–80.e8.

14. Xiu S, Dick A, Ju H, Mirzaie S, Abdi F, Cocklin S, et al. Inhibitors of SARS-CoV-2 Entry: Current and Future Opportunities. J Med Chem. 2020 Nov 12;63(21):12256–74.

15. Liu J, Cao R, Xu M, Wang X, Zhang H, Hu H, et al. Hydroxychloroquine, a less toxic derivative of chloroquine, is effective in inhibiting SARS-CoV-2 infection in vitro. Cell Discov. 2020 Mar 18;6:16.

16. Ou T, Mou H, Zhang L, Ojha A, Choe H, Farzan M. Hydroxychloroquine-mediated inhibition of SARS-CoV-2 entry is attenuated by TMPRSS2. PLoS Pathog. 2021 Jan;17(1):e1009212.

17. Savarino A. Expanding the frontiers of existing antiviral drugs: possible effects of HIV-1 protease inhibitors against SARS and avian influenza. J Clin Virol. 2005 Nov;34(3):170–8.

18. Mahdi M, Mótyán JA, Szojka ZI, Golda M, Miczi M, Tőzsér J. Analysis of the efficacy of HIV protease inhibitors against SARS-CoV-2’s main protease. Virol J. 2020 Nov 26;17(1):190.

19. Cao B, Wang Y, Wen D, Liu W, Wang J, Fan G, et al. A Trial of Lopinavir-Ritonavir in Adults Hospitalized with Severe Covid-19. N Engl J Med. 2020 May 7;382(19):1787–99.

20. Chen J, Xia L, Liu L, Xu Q, Ling Y, Huang D, et al. Antiviral Activity and Safety of Darunavir/Cobicistat for the Treatment of COVID-19. Open Forum Infect Dis. 2020 Jul;7(7):ofaa241.

21. Kim EJ, Choi SH, Park JS, Kwon YS, Lee J, Kim Y, et al. Use of Darunavir-Cobicistat as a Treatment Option for Critically Ill Patients with SARS-CoV-2 Infection. Yonsei Med J. 2020 Sep;61(9):826–30.

22. Jin Z, Du X, Xu Y, Deng Y, Liu M, Zhao Y, et al. Structure of M pro from SARS-CoV-2 and discovery of its inhibitors. Nature. 2020 Apr 9;582(7811):289–93.

23. Razzaghi-Asl N, Ebadi A, Shahabipour S, Gholamin D. Identification of a potential SARS-CoV2 inhibitor via molecular dynamics simulations and amino acid decomposition analysis. J Biomol Struct Dyn. 2020 Jul 24;1–16.

24. Mulangu S, Dodd LE, Davey RT Jr, Tshiani Mbaya O, Proschan M, Mukadi D, et al. A Randomized, Controlled Trial of Ebola Virus Disease Therapeutics. N Engl J Med. 2019 Dec 12;381(24):2293–303.

25. Jácome R, Campillo-Balderas JA, Ponce de León S, Becerra A, Lazcano A. Sofosbuvir as a potential alternative to treat the SARS-CoV-2 epidemic. Sci Rep. 2020 Jun 9;10(1):9294.

26. Chien M, Anderson TK, Jockusch S, Tao C, Li X, Kumar S, et al. Nucleotide Analogues as Inhibitors of SARS-CoV-2 Polymerase, a Key Drug Target for COVID-19. J Proteome Res. 2020 Nov 6;19(11):4690–7.

27. Jockusch S, Tao C, Li X, Anderson TK, Chien M, Kumar S, et al. A library of nucleotide analogues terminate RNA synthesis catalyzed by polymerases of coronaviruses that cause SARS and COVID-19. Antiviral Res. 2020 Aug;180:104857.

28. Cox RM, Wolf JD, Plemper RK. Therapeutically administered ribonucleoside analogue MK-4482/EIDD-2801 blocks SARS-CoV-2 transmission in ferrets. Nat Microbiol. 2021 Jan;6(1):11–8.

29. Kaptein SJF, Jacobs S, Langendries L, Seldeslachts L, ter Horst S, Liesenborghs L, et al. Favipiravir at high doses has potent antiviral activity in SARS-CoV-2−infected hamsters, whereas hydroxychloroquine lacks activity [Internet]. Vol. 117, Proceedings of the National Academy of Sciences. 2020. p. 26955–65. Available from: http://dx.doi.org/10.1073/pnas.2014441117

30. Klein S, Cortese M, Winter SL, Wachsmuth-Melm M, Neufeldt CJ, Cerikan B, et al. SARS-CoV-2 structure and replication characterized by in situ cryo-electron tomography. Nat Commun. 2020 Nov 18;11(1):5885.

31. Gupta MK, Vemula S, Donde R, Gouda G, Behera L, Vadde R. approaches to detect inhibitors of the human severe acute respiratory syndrome coronavirus envelope protein ion channel. J Biomol Struct Dyn. 2020 Apr 15;1–11.

32. Bobrowski T, Chen L, Eastman RT, Itkin Z, Shinn P, Chen CZ, et al. Synergistic and Antagonistic Drug Combinations against SARS-CoV-2. Mol Ther. 2021 Feb 3;29(2):873–85.

33. Leegwater E, Strik A, Wilms EB, Bosma LBE, Burger DM, Ottens TH, et al. Drug-induced Liver Injury in a Patient With Coronavirus Disease 2019: Potential Interaction of Remdesivir With P-Glycoprotein Inhibitors [Internet]. Clinical Infectious Diseases. 2020. Available from: http://dx.doi.org/10.1093/cid/ciaa883

34. Arribas JR, Arribas JR, Sanyal AJ, Soriano A, Chin BS, Chin BS, et al. 557. Impact of Concomitant Hydroxychloroquine Use on Safety and Efficacy of Remdesivir in Moderate COVID-19 Patients [Internet]. Vol. 7, Open Forum Infectious Diseases. 2020. p. S343–4. Available from: http://dx.doi.org/10.1093/ofid/ofaa439.751

35. Siegel D, Hui HC, Doerffler E, Clarke MO, Chun K, Zhang L, et al. Discovery and Synthesis of a Phosphoramidate Prodrug of a Pyrrolo[2,1-f][triazin-4-amino] Adenine C-Nucleoside (GS-5734) for the Treatment of Ebola and Emerging Viruses [Internet]. Vol. 60, Journal of Medicinal Chemistry. 2017. p. 1648–61. Available from: http://dx.doi.org/10.1021/acs.jmedchem.6b01594

36. Josephson F. Drug-drug interactions in the treatment of HIV infection: focus on pharmacokinetic enhancement through CYP3A inhibition. J Intern Med. 2010 Dec;268(6):530–9.

37. Deeks ED. Cobicistat: a review of its use as a pharmacokinetic enhancer of atazanavir and darunavir in patients with HIV-1 infection. Drugs. 2014 Feb;74(2):195–206.

38. Zanger UM, Schwab M. Cytochrome P450 enzymes in drug metabolism: regulation of gene expression, enzyme activities, and impact of genetic variation. Pharmacol Ther. 2013 Apr;138(1):103–41.

39. Wessler JD, Grip LT, Mendell J, Giugliano RP. The P-glycoprotein transport system and cardiovascular drugs. J Am Coll Cardiol. 2013 Jun 25;61(25):2495–502.

40. Sherman EM, Worley MV, Unger NR, Gauthier TP, Schafer JJ. Cobicistat: Review of a Pharmacokinetic Enhancer for HIV Infection. Clin Ther. 2015 Sep 1;37(9):1876–93.

41. Yamamoto N, Yang R, Yoshinaka Y, Amari S, Nakano T, Cinatl J, et al. HIV protease inhibitor nelfinavir inhibits replication of SARS-associated coronavirus. Biochem Biophys Res Commun. 2004 Jun 4;318(3):719–25.

42. Yamamoto N, Matsuyama S, Hoshino T, Yamamoto N. Nelfinavir inhibits replication of severe acute respiratory syndrome coronavirus 2 in vitro [Internet]. Available from: http://dx.doi.org/10.1101/2020.04.06.026476

43. Sharma P, Vijayan V, Pant P, Sharma M, Vikram N, Kaur P, et al. Identification of potential drug candidates to combat COVID-19: a structural study using the main protease (mpro) of SARS-CoV-2. J Biomol Struct Dyn. 2020 Aug 3;1–11.

44. Ibrahim MAA, Abdelrahman AHM, Hegazy M-EF. drug repurposing and molecular dynamics puzzled out potential SARS-CoV-2 main protease inhibitors. J Biomol Struct Dyn. 2020 Jul 20;1– 12.

45. Ahmed S, Mahtarin R, Ahmed SS, Akter S, Islam MS, Mamun AA, et al. Investigating the binding affinity, interaction, and structure-activity-relationship of 76 prescription antiviral drugs targeting RdRp and Mpro of SARS-CoV-2. J Biomol Struct Dyn. 2020 Jul 28;1–16.

46. Mathias AA, German P, Murray BP, Wei L, Jain A, West S, et al. Pharmacokinetics and pharmacodynamics of GS-9350: a novel pharmacokinetic enhancer without anti-HIV activity. Clin Pharmacol Ther. 2010 Mar;87(3):322–9.

47. Zhang L, Lin D, Sun X, Curth U, Drosten C, Sauerhering L, et al. Crystal structure of SARS-CoV-2 main protease provides a basis for design of improved α-ketoamide inhibitors. Science. 2020 Apr 24;368(6489):409–12.

48. Huang K, Luo S, Cong Y, Zhong S, Zhang JZH, Duan L. An accurate free energy estimator: based on MM/PBSA combined with interaction entropy for protein-ligand binding affinity. Nanoscale. 2020 May 21;12(19):10737–50.

49. Pape C, Remme R, Wolny A, Olberg S, Wolf S, Cerrone L, et al. Microscopy-based assay for semi-quantitative detection of SARS-CoV-2 specific antibodies in human sera [Internet]. BioEssays. 2020. p. 2000257. Available from: http://dx.doi.org/10.1002/bies.202000257

50. Ou X, Liu Y, Lei X, Li P, Mi D, Ren L, et al. Characterization of spike glycoprotein of SARS-CoV-2 on virus entry and its immune cross-reactivity with SARS-CoV. Nat Commun. 2020 Mar 27;11(1):1620.

51. Algaissi A, Alfaleh MA, Hala S, Abujamel TS, Alamri SS, Almahboub SA, et al. SARS-CoV-2 S1 and N-based serological assays reveal rapid seroconversion and induction of specific antibody response in COVID-19 patients. Sci Rep. 2020 Oct 6;10(1):16561.

52. Watanabe R, Matsuyama S, Shirato K, Maejima M, Fukushi S, Morikawa S, et al. Entry from the cell surface of severe acute respiratory syndrome coronavirus with cleaved S protein as revealed by pseudotype virus bearing cleaved S protein. J Virol. 2008 Dec;82(23):11985–91.

53. Yin W, Mao C, Luan X, Shen D-D, Shen Q, Su H, et al. Structural basis for inhibition of the RNA-dependent RNA polymerase from SARS-CoV-2 by remdesivir. Science. 2020 Jun 26;368(6498):1499–504.

54. Daina A, Michielin O, Zoete V. SwissADME: a free web tool to evaluate pharmacokinetics, drug-likeness and medicinal chemistry friendliness of small molecules. Sci Rep. 2017 Mar 3;7:42717.

55. Pires DEV, Blundell TL, Ascher DB. pkCSM: Predicting Small-Molecule Pharmacokinetic and Toxicity Properties Using Graph-Based Signatures. J Med Chem. 2015 May 14;58(9):4066–72.

56. Tian S, Djoumbou-Feunang Y, Greiner R, Wishart DS. CypReact: A Software Tool for in Silico Reactant Prediction for Human Cytochrome P450 Enzymes. J Chem Inf Model. 2018 Jun 25;58(6):1282–91.

57. von Richter O, Burk O, Fromm MF, Thon KP, Eichelbaum M, Kivistö KT. Cytochrome P450 3A4 and P-glycoprotein expression in human small intestinal enterocytes and hepatocytes: a comparative analysis in paired tissue specimens. Clin Pharmacol Ther. 2004 Mar;75(3):172–83.

58. Bradley G, Ling V. P-glycoprotein, multidrug resistance and tumor progression. Cancer Metastasis Rev. 1994 Jun;13(2):223–33.

59. Stanifer ML, Kee C, Cortese M, Zumaran CM, Triana S, Mukenhirn M, et al. Critical Role of Type III Interferon in Controlling SARS-CoV-2 Infection in Human Intestinal Epithelial Cells. Cell Rep. 2020 Jul 7;32(1):107863.

60. Bartlett JA, DeMasi R, Quinn J, Moxham C, Rousseau F. Overview of the effectiveness of triple combination therapy in antiretroviral-naive HIV-1 infected adults. AIDS. 2001 Jul 27;15(11):1369– 77.

61. Naggie S, Muir AJ. Oral Combination Therapies for Hepatitis C Virus Infection: Successes, Challenges, and Unmet Needs. Annu Rev Med. 2017 Jan 14;68:345–58.

62. van Waterschoot RAB, ter Heine R, Wagenaar E, van der Kruijssen CMM, Rooswinkel RW, Huitema ADR, et al. Effects of cytochrome P450 3A (CYP3A) and the drug transporters P-glycoprotein (MDR1/ABCB1) and MRP2 (ABCC2) on the pharmacokinetics of lopinavir. Br J Pharmacol. 2010 Jul;160(5):1224–33.

63. White KM, Rosales R, Yildiz S, Kehrer T, Miorin L, Moreno E, et al. Plitidepsin has potent preclinical efficacy against SARS-CoV-2 by targeting the host protein eEF1A. Science [Internet]. 2021 Jan 25; Available from: http://dx.doi.org/10.1126/science.abf4058

64. Brandon EFA, Sparidans RW, van Ooijen RD, Meijerman I, Lazaro LL, Manzanares I, et al. In vitro characterization of the human biotransformation pathways of aplidine, a novel marine anti-cancer drug. Invest New Drugs. 2007 Feb;25(1):9–19.

65. Kim K-A, Park J-Y, Lee J-S, Lim S. Cytochrome P450 2C8 and CYP3A4/5 are involved in chloroquine metabolism in human liver microsomes. Arch Pharm Res. 2003 Aug;26(8):631–7.

66. Hu W-J, Chang L, Yang Y, Wang X, Xie Y-C, Shen J-S, et al. Pharmacokinetics and tissue distribution of remdesivir and its metabolites nucleotide monophosphate, nucleotide triphosphate, and nucleoside in mice. Acta Pharmacol Sin [Internet]. 2020 Oct 12; Available from: http://dx.doi.org/10.1038/s41401-020-00537-9

67. Lepist E-I, Phan TK, Roy A, Tong L, Maclennan K, Murray B, et al. Cobicistat boosts the intestinal absorption of transport substrates, including HIV protease inhibitors and GS-7340, in vitro. Antimicrob Agents Chemother. 2012 Oct;56(10):5409–13.

68. Jorgensen SCJ, Kebriaei R, Dresser LD. Remdesivir: Review of Pharmacology, Pre-clinical Data, and Emerging Clinical Experience for COVID-19. Pharmacotherapy. 2020 Jul;40(7):659–71.

69. Hoffmann M, Kleine-Weber H, Pöhlmann S. A Multibasic Cleavage Site in the Spike Protein of SARS-CoV-2 Is Essential for Infection of Human Lung Cells. Mol Cell. 2020 May 21;78(4):779– 84.e5.

70. Yang K. What Do We Know About Remdesivir Drug Interactions? Clin Transl Sci. 2020 Sep;13(5):842–4.

71. Wang P, Shehu AI, Liu K, Lu J, Ma X. Biotransformation of Cobicistat: Metabolic Pathways and Enzymes. Drug Metab Lett. 2016;10(2):111–23.

72. Wishart DS, Feunang YD, Guo AC, Lo EJ, Marcu A, Grant JR, et al. DrugBank 5.0: a major update to the DrugBank database for 2018. Nucleic Acids Res. 2018 Jan 4;46(D1):D1074–82.

73. Andrianov AM, Kornoushenko YV, Karpenko AD, Bosko IP, Tuzikov AV. Computational discovery of small drug-like compounds as potential inhibitors of SARS-CoV-2 main protease. J Biomol Struct Dyn. 2020 Jul 14;1–13.

74. Friesner RA, Banks JL, Murphy RB, Halgren TA, Klicic JJ, Mainz DT, et al. Glide: a new approach for rapid, accurate docking and scoring. 1. Method and assessment of docking accuracy. J Med Chem. 2004 Mar 25;47(7):1739–49.

75. Maier JA, Martinez C, Kasavajhala K, Wickstrom L, Hauser KE, Simmerling C. ff14SB: Improving the Accuracy of Protein Side Chain and Backbone Parameters from ff99SB. J Chem Theory Comput. 2015 Aug 11;11(8):3696–713.

76. Wang J, Wolf RM, Caldwell JW, Kollman PA, Case DA. Development and testing of a general amber force field. J Comput Chem. 2004 Jul 15;25(9):1157–74.

77. Roe DR, Cheatham TE 3rd. PTRAJ and CPPTRAJ: Software for Processing and Analysis of Molecular Dynamics Trajectory Data. J Chem Theory Comput. 2013 Jul 9;9(7):3084–95.

78. Miller BR 3rd, McGee TD Jr, Swails JM, Homeyer N, Gohlke H, Roitberg AE. MMPBSA.py: An Efficient Program for End-State Free Energy Calculations. J Chem Theory Comput. 2012 Sep 11;8(9):3314–21.

79. Duan L, Liu X, Zhang JZH. Interaction Entropy: A New Paradigm for Highly Efficient and Reliable Computation of Protein-Ligand Binding Free Energy. J Am Chem Soc. 2016 May 4;138(17):5722– 8.

80. Cortese M, Lee J-Y, Cerikan B, Neufeldt CJ, Oorschot VMJ, Köhrer S, et al. Integrative Imaging Reveals SARS-CoV-2-Induced Reshaping of Subcellular Morphologies. Cell Host Microbe. 2020 Dec 9;28(6):853–66.e5.

81. Fischl W, Bartenschlager R. High-throughput screening using dengue virus reporter genomes. Methods Mol Biol. 2013;1030:205–19.

82. Livak KJ, Schmittgen TD. Analysis of relative gene expression data using real-time quantitative PCR and the 2(-Delta Delta C(T)) Method. Methods. 2001 Dec;25(4):402–8.

83. Shytaj IL, Lucic B, Forcato M, Penzo C, Billingsley J, Laketa V, et al. Alterations of redox and iron metabolism accompany the development of HIV latency. EMBO J. 2020 May 4;39(9):e102209.

84. Feoktistova M, Geserick P, Leverkus M. Crystal Violet Assay for Determining Viability of Cultured Cells. Cold Spring Harb Protoc. 2016 Apr 1;2016(4):db.prot087379.

85. Schindelin J, Arganda-Carreras I, Frise E, Kaynig V, Longair M, Pietzsch T, et al. Fiji: an open-source platform for biological-image analysis. Nat Methods. 2012 Jun 28;9(7):676–82.

86. Ianevski A, He L, Aittokallio T, Tang J. SynergyFinder: a web application for analyzing drug combination dose-response matrix data. Bioinformatics. 2020 Apr 15;36(8):2645.

87. Yadav B, Wennerberg K, Aittokallio T, Tang J. Searching for Drug Synergy in Complex Dose-Response Landscapes Using an Interaction Potency Model. Comput Struct Biotechnol J. 2015 Sep 25;13:504–13.

88. Kakuda TN, Van De Casteele T, Petrovic R, Neujens M, Salih H, Opsomer M, et al. Bioequivalence of a darunavir/cobicistat fixed-dose combination tablet versus single agents and food effect in healthy volunteers. Antivir Ther. 2014 Jun 25;19(6):597–606.

89. Ma C, Sacco MD, Hurst B, Townsend JA, Hu Y, Szeto T, et al. Boceprevir, GC-376, and calpain inhibitors II, XII inhibit SARS-CoV-2 viral replication by targeting the viral main protease. Cell Res. 2020 Aug;30(8):678–92.

90. Chou T-C. Drug combination studies and their synergy quantification using the Chou-Talalay method. Cancer Res. 2010 Jan 15;70(2):440–6.

91. Korolev D, Balakin KV, Nikolsky Y, Kirillov E, Ivanenkov YA, Savchuk NP, et al. Modeling of human cytochrome p450-mediated drug metabolism using unsupervised machine learning approach. J Med Chem. 2003 Aug 14;46(17):3631–43.

92. Yap CW, Chen YZ. Prediction of cytochrome P450 3A4, 2D6, and 2C9 inhibitors and substrates by using support vector machines. J Chem Inf Model. 2005 Jul;45(4):982–92.

93. Terfloth L, Bienfait B, Gasteiger J. Ligand-based models for the isoform specificity of cytochrome P450 3A4, 2D6, and 2C9 substrates. J Chem Inf Model. 2007 Jul;47(4):1688–701.

94. Michielan L, Terfloth L, Gasteiger J, Moro S. Comparison of multilabel and single-label classification applied to the prediction of the isoform specificity of cytochrome p450 substrates. J Chem Inf Model. 2009 Nov;49(11):2588–605.

95. Ramesh M, Bharatam PV. CYP isoform specificity toward drug metabolism: analysis using common feature hypothesis. J Mol Model. 2012 Feb;18(2):709–20.

96. Nembri S, Grisoni F, Consonni V, Todeschini R. In Silico Prediction of Cytochrome P450-Drug Interaction: QSARs for CYP3A4 and CYP2C9. Int J Mol Sci [Internet]. 2016 Jun 9;17(6). Available from: http://dx.doi.org/10.3390/ijms17060914

97. Zhang T, Dai H, Liu LA, Lewis DFV, Wei D. Classification Models for Predicting Cytochrome P450 Enzyme-Substrate Selectivity. Mol Inform. 2012 Jan;31(1):53–62.

98. Yamashita F, Hara H, Ito T, Hashida M. Novel hierarchical classification and visualization method for multiobjective optimization of drug properties: application to structure-activity relationship analysis of cytochrome P450 metabolism. J Chem Inf Model. 2008 Feb;48(2):364–9.

