## Supplementary figures and images for "The FDA-approved drug cobicistat synergizes with remdesivir to inhibit SARS-CoV-2 replication"

### Supplementary Figure 1

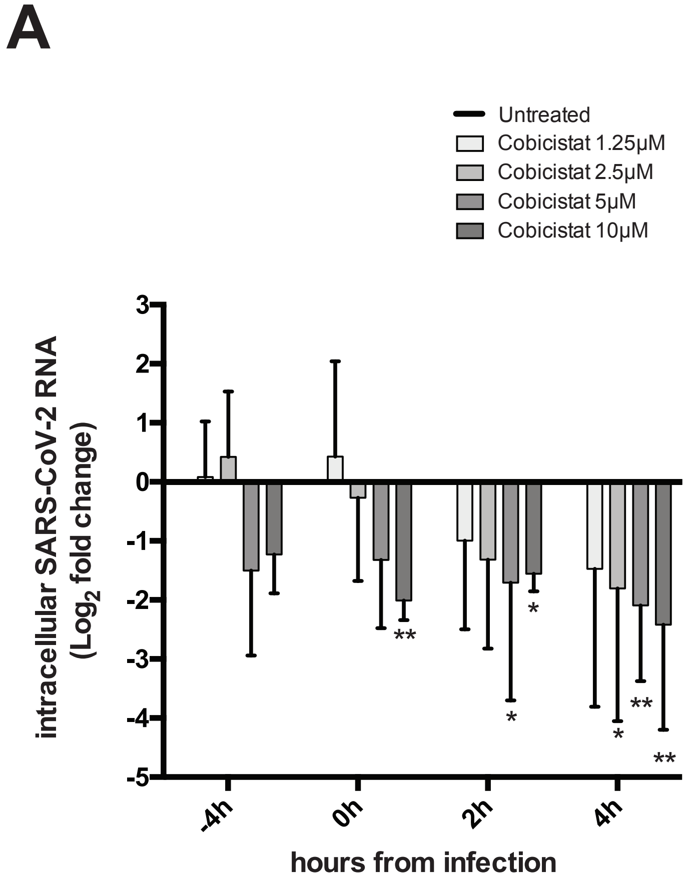


intracellular

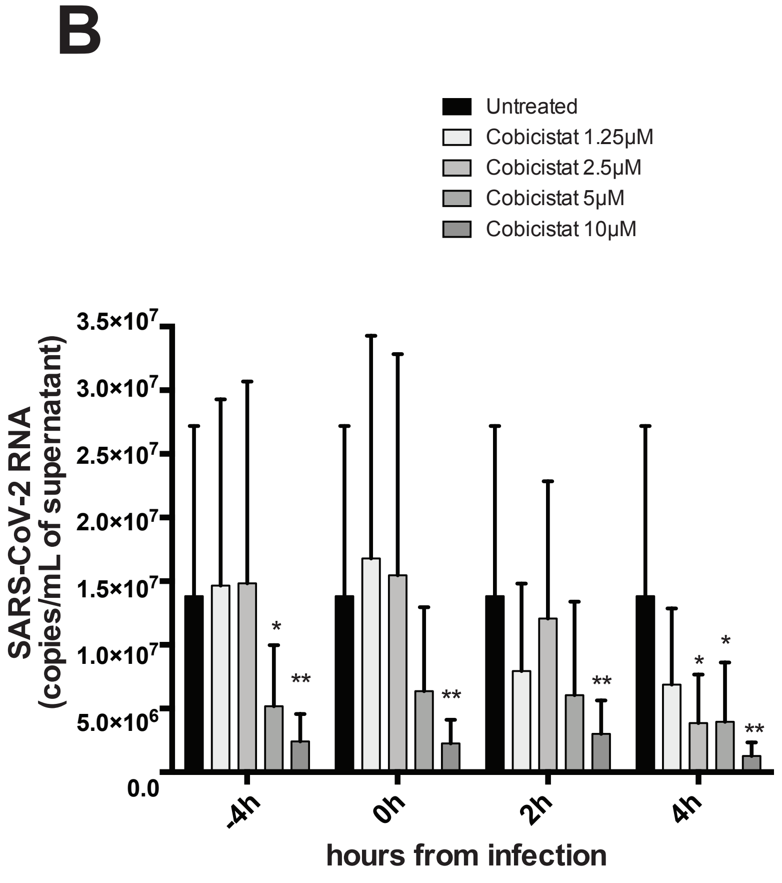


supernatant

### Supplementary Figure 2

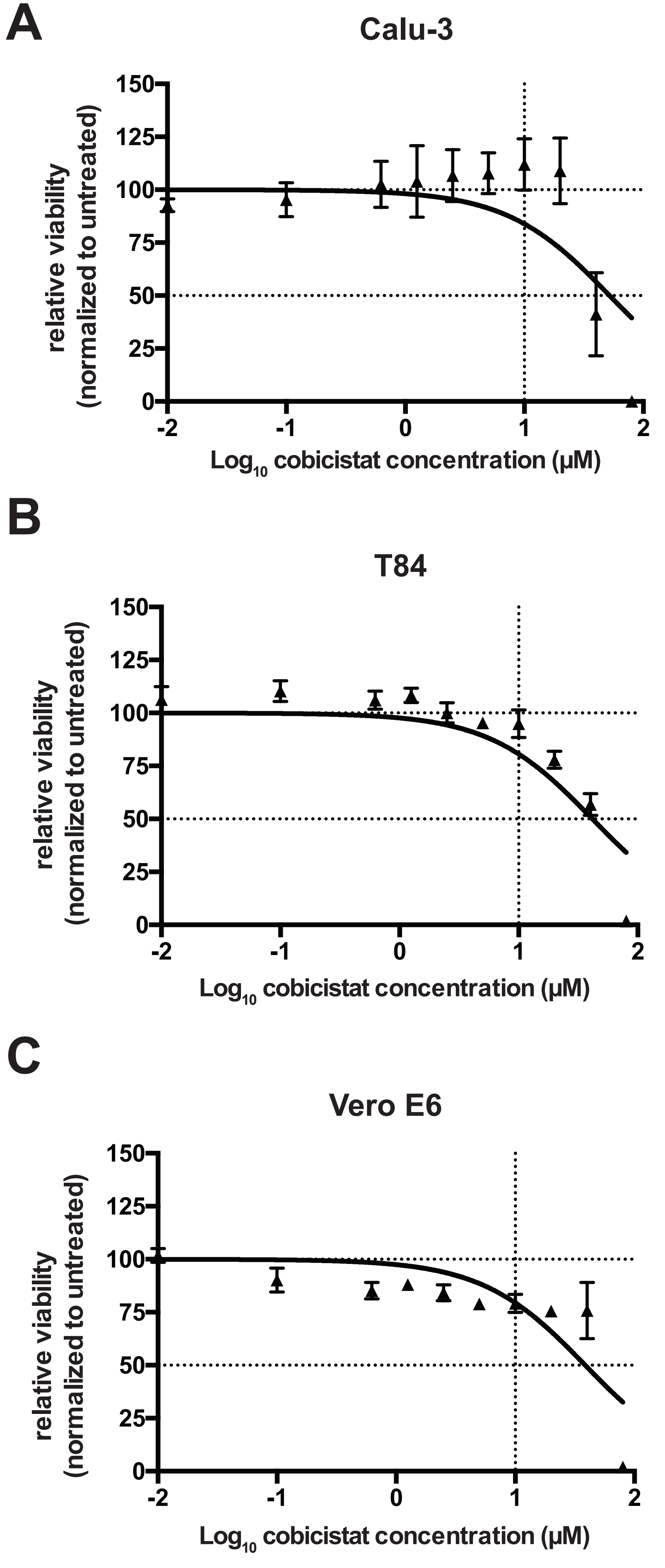

### Supplementary Figure 3

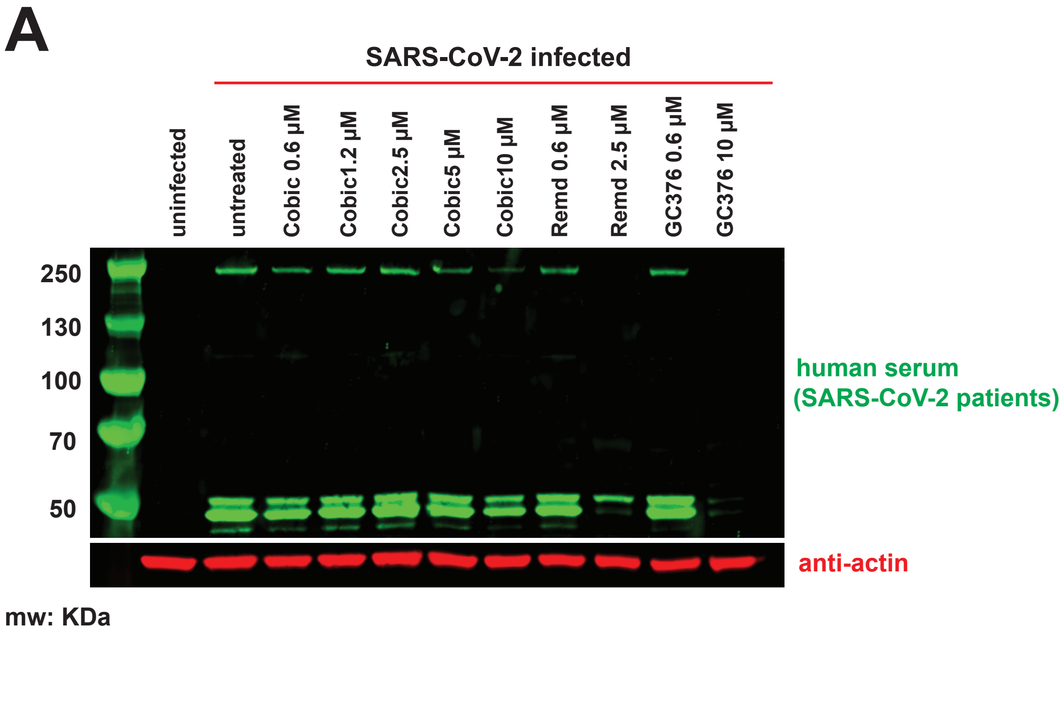

### Supplementary Figure 4

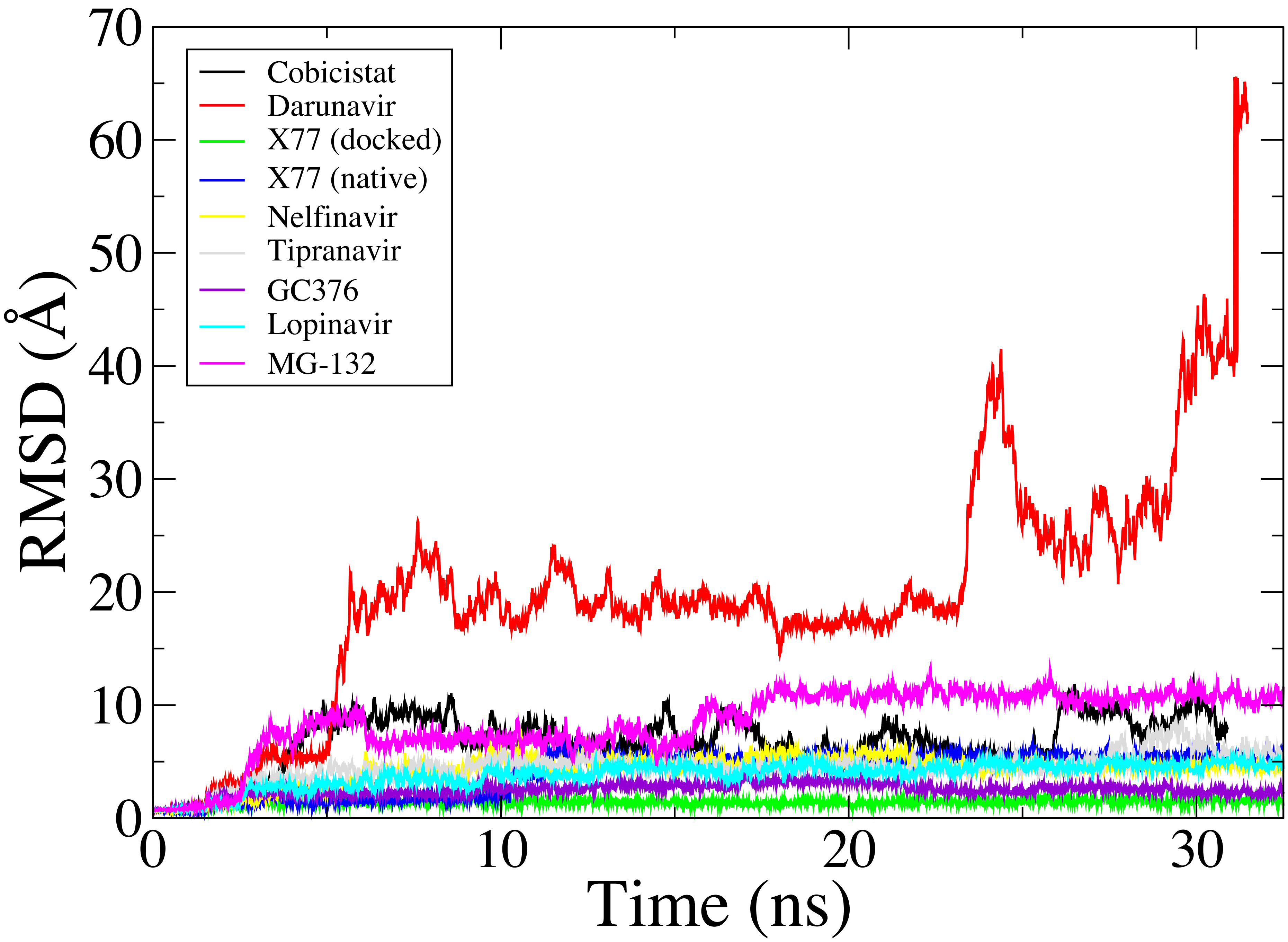

### Supplementary Figure 5

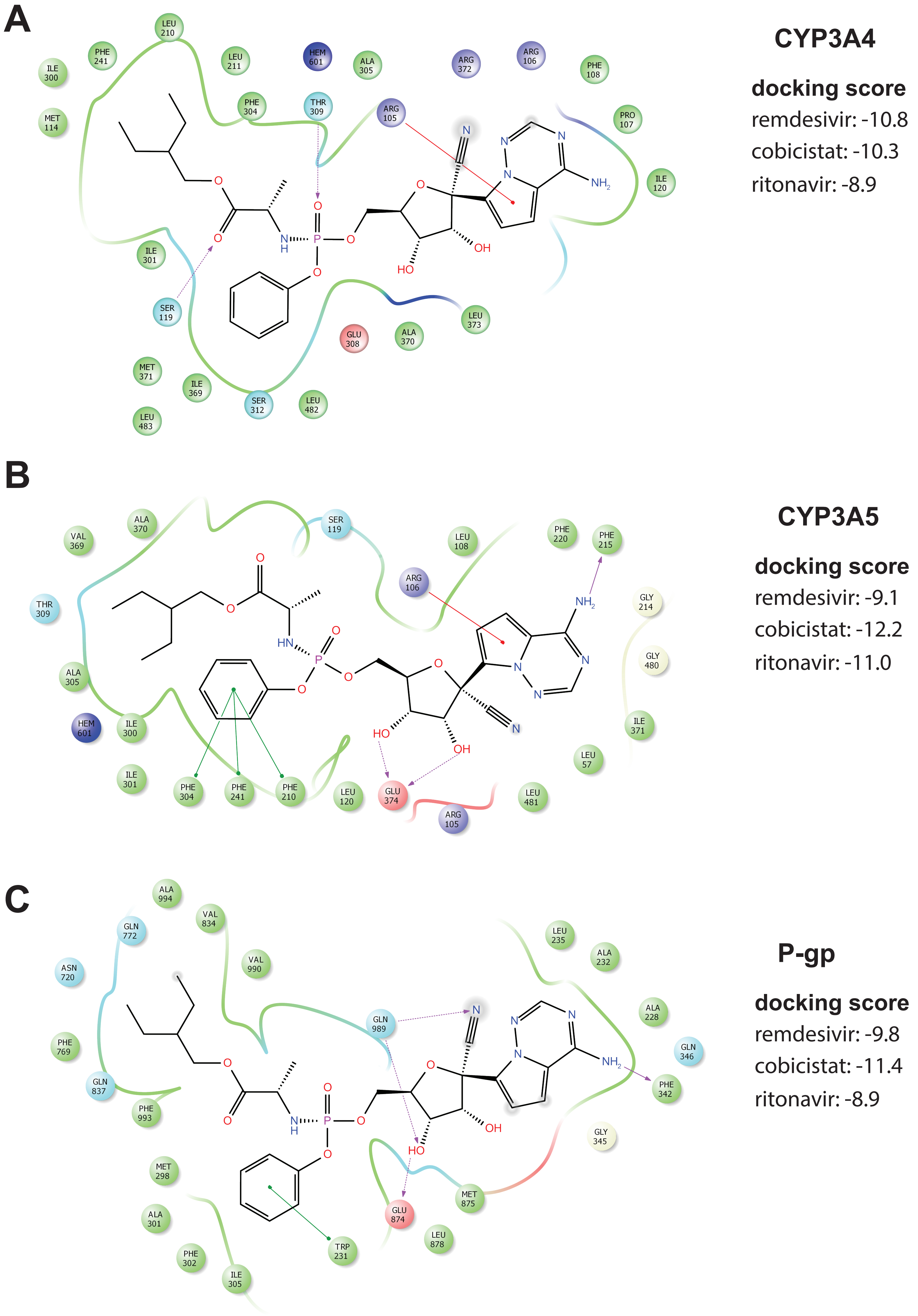

### Supplementary Figure 6

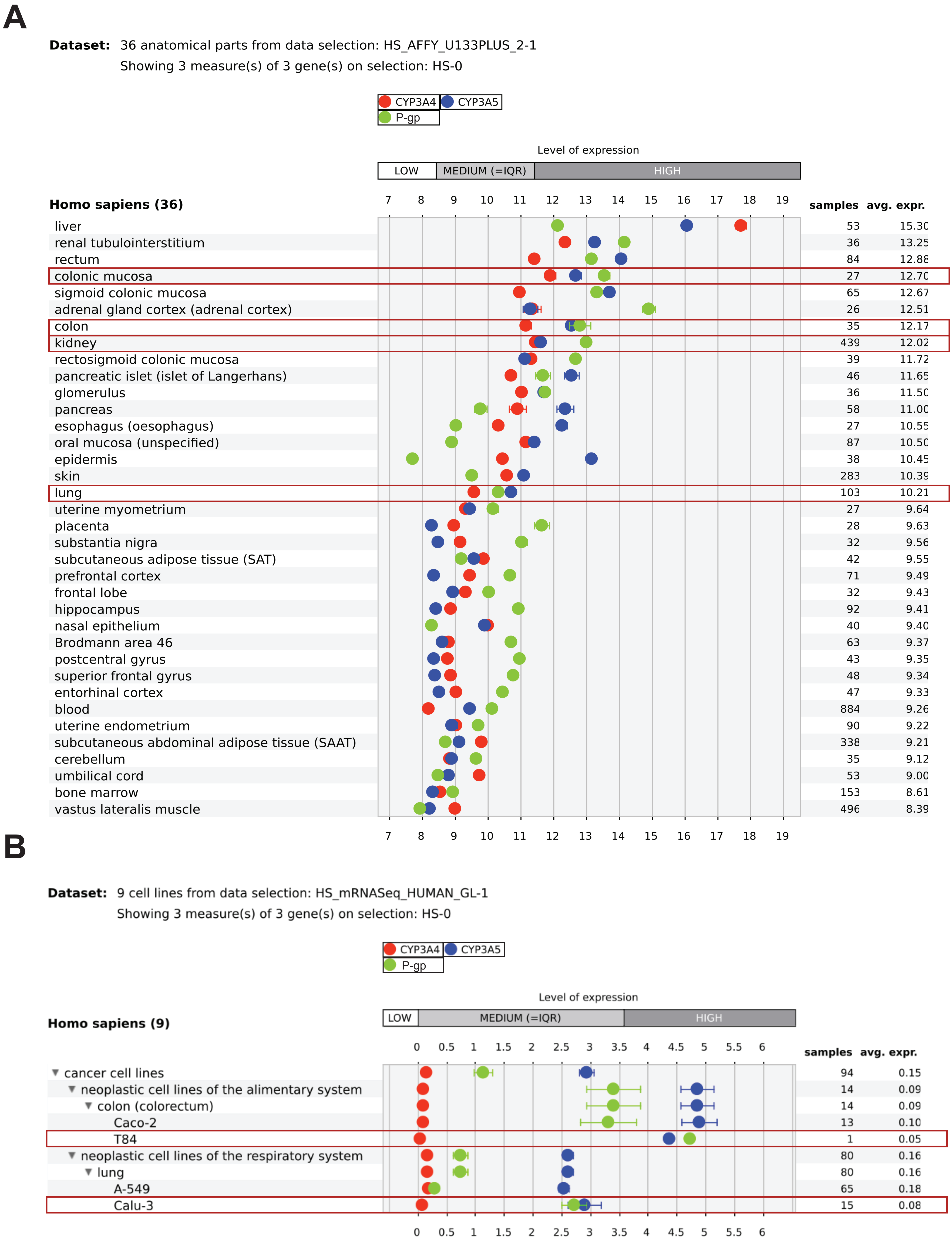

### Supplementary Figure 7

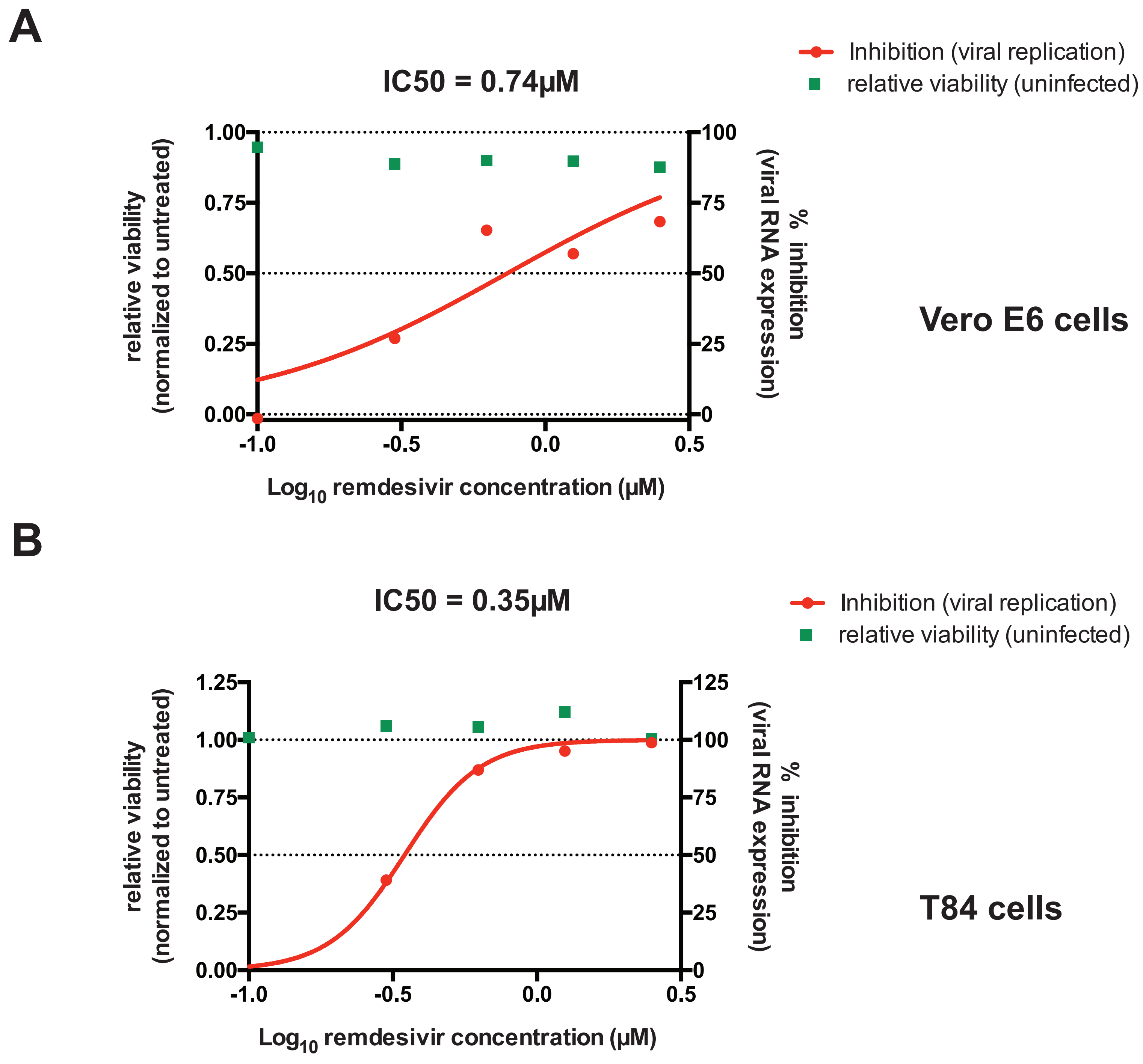

### Supplementary Figure 8

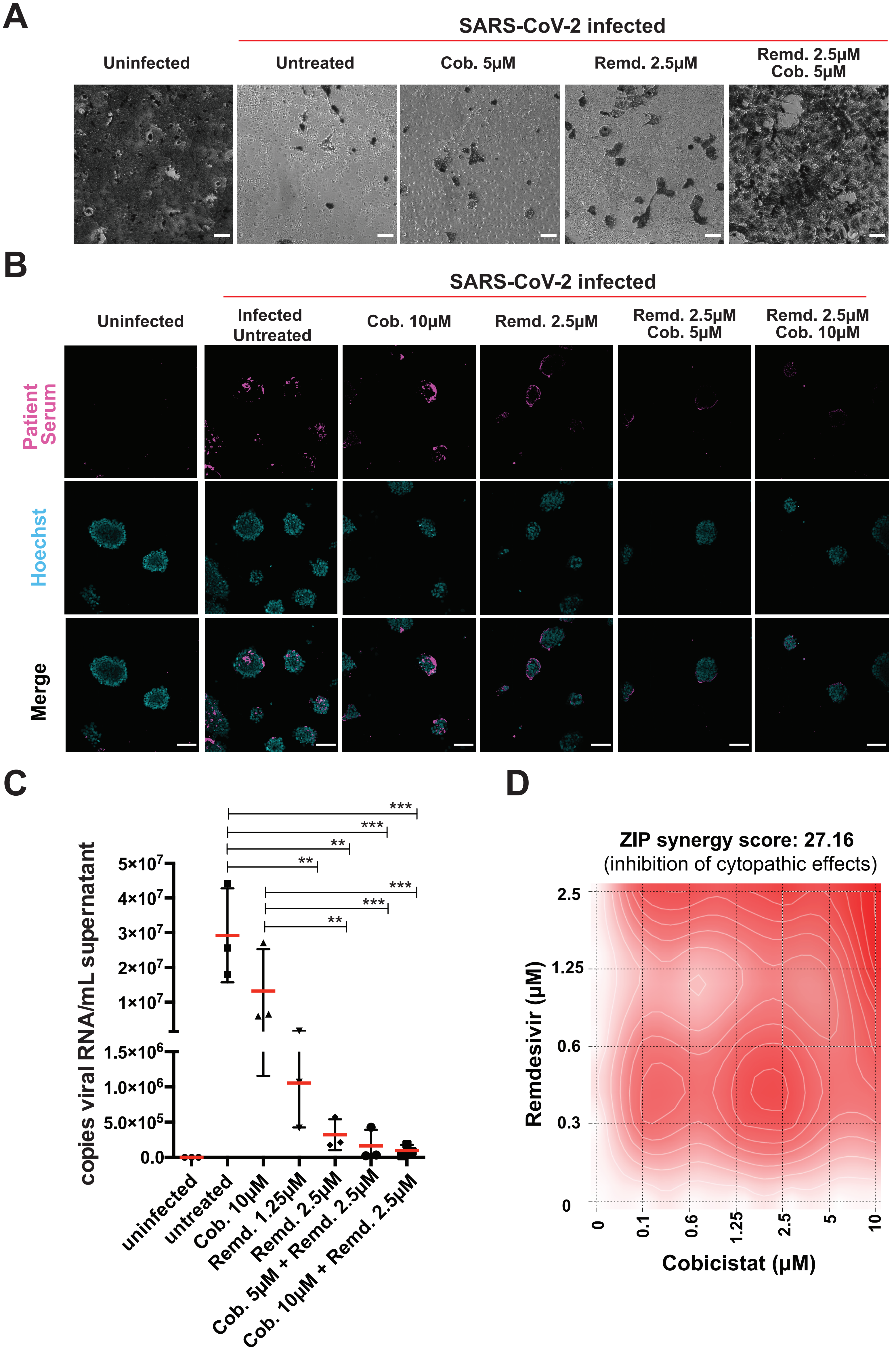
