## Supplementary Table 1 for "The FDA-approved drug cobicistat synergizes with remdesivir to inhibit SARS-CoV-2 replication"

| DRUGBANK_ID | Drug groups | Generic name | Main indication | Docking score |
|---|---|---|---|---|
| DB01362 | approved | Iohexol | Contrast agent | -11.72 |
| DB09134 | approved | Ioversol | Contrast agent | -11.03 |
| DB12407 | approved; investigational | Iobitridol | Contrast agent | -10.22 |
| DB12615 | approved; investigational | Plazomicin | Antibiotic for urinary tract infections | -9.43 |
| DB00932 | approved; investigational | Tipranavir | HIV protease inhibitor | -8.06 |
| DB00220 | approved | Nelfinavir | HIV protease inhibitor | -7.91 |
| DB08909 | approved | Glycerol phenylbutyrate | Nitrogen-binding agent for management of urea cycle disorders | -7.86 |
| DB00905 | approved; investigational | Bimatoprost | Analog of prostaglandin F2α for treatment of glaucoma | -7.67 |
| DB08889 | approved; investigational | Carfilzomib | Proteasome Inhibitor (anti-cancer) | -7.54 |
| **DB09065** | **approved** | **Cobicistat** | **CYP3A inhibitor for boosting HIV-1 protease inhibitors** | **-7.12** |
| DB04868 | approved; investigational | Nilotinib | Tyrosine kinase inhibitor for treatment of chronic myelogenous leukemia | -7.05 |
| DB01288 | approved; investigational | Fenoterol | Beta adrenergic agonist for asthma treatment | -7.05 |
| DB00482 | approved; investigational | Celecoxib | Nonsteroidal anti-inflammatory drug | -6.80 |
| DB13931 | approved | Netarsudil | Rho kinase inhibitor for treatment of glaucoma | -6.75 |
| DB11611 | approved | Lifitegrast | Anti-Inflammatory for treatment of keratoconjunctivitis sicca | -6.45 |
| DB11979 | approved; investigational | Elagolix | gonadotropin-releasing hormone antagonist for treatment of endometriosis pain | -5.72 |
| DB01116 | approved; investigational | Trimethaphan | nicotinic antagonist used to counteract hypertension | -5.70 |
