## Supplementary Table 2 for "The FDA-approved drug cobicistat synergizes with remdesivir to inhibit SARS-CoV-2 replication"

**List of qPCR primers used in the study**

| name | sequence | source |
|---|---|---|
| **2019-nCoV_N1-Forward** | GAC CCC AAA ATC AGC GAA AT | https://www.cdc.gov/coronavirus/2019-ncov/lab/rt-pcr-panel-primer-probes.html |
| **2019-nCoV_N1-Reverse** | TCT GGT TAC TGC CAG TTG AAT CTG | |
| **2019-nCoV_N2-Forward** | TTA CAA ACA TTG GCC GCA AA | https://www.cdc.gov/coronavirus/2019-ncov/lab/rt-pcr-panel-primer-probes.html |
| **2019-nCoV_N2-Reverse** | GCG CGA CAT TCC GAA GAA | |
| **Hum Cyp3A4-Forward** | TGA TGG CTC TCA TCC CAG AC | |
| **Cyp3A4-Reverse** | AGC CCC ACA CTT TTC CAT AC | |
| **AGM Cyp3A4-Forward** | TGA TGG ACC TCA TCC CAG AC | |
| **Hum Cyp3A5-Forward** | CGA CAA ACA AAA GCA CCG AC | |
| **Hum Cyp3A5-Reverse** | TTA TTG ACT GGG CTG CGA G | |
| **AGM Cyp3A5-Forward** | CGA CAA ACA AAA GCA CCG AG | |
| **AGM Cyp3A5-Reverse** | TAA TTG ATT GGG CCA CGA G | |
| **P-gp (MDR1)-F** | CCC ATC ATT GCA ATA GCA GG | Gao et al. Int J Clin Exp Pathol 2015 |
| **P-gp (MDR1)-R** | TGT TCA AAC TTC TGC TCC TGA | |
| **TBP-F** | CCA CTC ACA GAC TCT CAC AAC | Stanifer et al. Cell Reports 2020 |
| **TBP-R** | CTG CGG TAC AAT CCC AGA ACT | |

Hum = human
AGM = african green monkey (Vero E6 cells)
