## Supplementary Table 3 for "The FDA-approved drug cobicistat synergizes with remdesivir to inhibit SARS-CoV-2 replication"

| Ligand | ΔG solvation | ΔE interactions | -TΔS | ΔG total | FRET-determined EC50 (3CLpro) |
|---|---|---|---|---|---|
| GC376 | 35.9 | -69.3 | 18.2 | -15.2 | 0.11 µM |
| X77 (docked) | 51.1 | -88.9 | 25.6 | -12.2 | N.A. |
| X77 (native) | 34.4 | -62.8 | 17 | -11.4 | N.A. |
| Tipranavir | 28.8 | -54.4 | 19.5 | -6 | 47 µM |
| Lopinavir | 28.3 | -61.9 | 28.6 | -5 | 219 µM |
| MG-132 | 19.1 | -41.6 | 18 | -4.5 | 18 µM |
| Darunavir | 10.9 | -17.8 | 17.1 | 10.3 | could not be calculated |
| Cobicistat | 82.9 | -112.8 | 44 | 14.2 | could not be calculated |
| Nelfinavir | 112.7 | -152.4 | 81.3 | 41.6 | could not be calculated |
